# Type IV minor pilin ComN predicted the USS-receptor in Pasteurellaceae

**DOI:** 10.1101/2025.06.16.659825

**Authors:** Stian Aleksander Helsem, Kristian Alfsnes, Stephan A. Frye, Alexander Hesselberg Løvestad, Ole Herman Ambur

## Abstract

The Uptake Signal Sequence (USS) receptor, facilitating acquisition of homologous DNA by natural transformation in *Haemophilus influenzae* and other Pasteurellaceae, remains unknown. Assuming extracellular USS-binding in accordance with experimental data and existing models of the transformation process, discriminating functional gene ontology assessment, cellular localization predictions and deep-learning structural modeling of protein-DNA complexes, a prepilin peptidase-dependent protein A (PpdA), was identified as the strongest USS receptor candidate in different Pasteurellaceae family members with divergent USS specificities. PpdA was the only orthogroup to be modeled to form specific protein-USS complexes significantly better than complexes with sequence-scrambled versions of USS by AlphaFold3. Further analyses of PpdA complexes, from ten different Pasteurellaceae with divergent USS-type enrichment, using geometric deep learning protein-DNA sequence specificity predictions and coevolution analyses were found to further support this USS receptor candidacy. PpdA was found to share overall structural domains with the non-sequence specific DNA receptor FimT in *Legionella pneumophila* and a particular β-sheet transversing disulfide-bridge with ComP, the DNA uptake sequence receptor of the Neisseriaceae. In compliance with a previously given gene name, we propose ComN to be used for these PpdA orthologs which structurally and evolutionary were here predicted to be the USS-receptors in Pasteurellaceae.

## Introduction

Deep-learning protein structure prediction has revolutionized biological research in recent years, an impact recognized by the Nobel prize in Chemistry 2024. The development of new tools and benchmarking studies involving crystallographic detail has allowed the field to obtain better functional understanding from predicted 3D structures, although models cannot represent ground truths (discussed in (1)). Recently, tools that model interactions between proteins and ligands and DNA/RNA have been developed and applied to address specific biological phenomena. Three such tools, AlphaFold3, RosettaFold-2NA and Chai-1 have been developed to model protein-DNA complexes (2–4). The input data are simply protein and DNA linear amino acid and nucleotide sequences, respectively, which are folded in learned statistical pattern recognition algorithms. In our previous study, these three AI-tools were applied to study modeled complexes of DNA and the minor type IV pilin ComP, which is the DNA Uptake Sequence (DUS)-specific protein in the bacterial Neisseriaceae family (5). The protein responsible for the equivalent specific uptake process in the other known DNA uptake-specific family, the Pasteurellaceae, remains unknown. Several deep-learning algorithms have recently been developed to address a general need to go from protein-DNA complex structure modeling to specificity prediction. Deep Predictor of Binding Specificity (DeepPBS) is such and takes its approach by combining geometric convolution predictions of protein and DNA structures (6). DeepPBS input data can be either experimentally generated, simulated or predicted structures and this approach was cross-validated and benchmarked for a range of protein families (6). Natural transformation is the uptake and homologous recombination of DNA from the extracellular environment and is a feature of many bacterial species. The molecular mechanisms regulating and mediating bacterial transformation show great diversity and thereby signify the biological essence of allelic reshuffles through bacterial sex (7). A unifying feature of the transformation process in most Gram-negative and positive species are type IV pili (T4P) used to capture extracellular DNA (8). Current models show how DNA binds to the protruding pilus and is pulled onto the plasma membrane together with the depolymerizing and retracting pilus. Passage of DNA through the plasma membrane happens in single stranded form through the transmembrane ComEC-pore, conserved in both Gram groups. Once in the cytoplasm DNA is processed by DprA and homologous recombination is achieved by the means of RecA, a ubiquitous protein in nature. Extracellular DNA binding may be either specific or unspecific in different bacteria.

Neisseriaceae (ꞵ-proteobacteria) and Pasteurellaceae (γ-proteobacteria) are the only families known to discriminate homologous from heterologous DNA at this early step in the transformation process by specific binding to short (ca. 9-12 nt) DNA-motifs which are highly enriched in respective genomes (9–15). Notable human pathogens in these families are *Neisseria gonorrhoeae, Neisseria meningitidis* of the Neisseriaceae and *H. influenzae* and *Pasteurella multocida* of the Pasteurellaceae. The physiological state of competence for transformation is constitutive in the *Neisseria*, while it is induced by cyclic AMP (a starvation and lag-phase signal) in the Pasteurellaceae (16,17). The induction of expression of seventeen *H. influenzae* genes has been shown to be required for transformation and the gene/operon regulatory pathway (Sxy) has been characterized (18). The short DNA motifs recognized by Pasteurellaceae and Neisseriaceae are by tradition named Uptake Signal Sequences (USS) and DNA Uptake Sequences (DUS), respectively. USS and DUS are highly dissimilar in sequence motif and are therefore considered to have evolved convergently to achieve the same means: bias transformation to involve homologous DNA (14).

Characteristic USS/DUS containing genomes harbour hundreds or thousands of these motifs which are often organized as transcriptional terminators, yet the presence of a single motif is sufficient to increase transformation rates (9). The DUS and USS motifs have both diverged into distinct dialects/variants or types within each bacterial family and may differ by 1-3 nucleotides from each other, yet with their respective transformation and uptake-essential inner cores conserved. The inner core sequences are 5’-CTG-3’ in the Neisseriaceae DUS and 5’-GCGG-3’ in Pasteurellaceae USS (19,20). A total of eight dialects have been described in the Neisseriaceae (AT-DUS, AG-DUS, AG-mucDUS, AG-eikDUS, AG-kingDUS, AA-king3 DUS, TG-wadDUS and AG-simDUS) and two USS-types in the Pasteurellaceae classified as *Haemophilus*-type Hin-USS (5’-AAGTGCGGT-3’) and *Actinobacillus pleuropneumoniae* (now *Aggregatibacter pleuropneumoniae*)-type Apl-USS (5’-ACAAGCGGT-3’) (19,21).

Both DUS and USS dialects adhere to phylogeny with genomic enrichment within defined subclades (19,21). DUS are confined to continuous motifs extending up to about 13 nucleotides in length whereas the USS are discontinuous and are characterized by the 9-10 nt core motifs and two less conserved and discontinuous downstream AT-rich regions making the USS motifs extend up to 32 nucleotides (21). These AT-rich regions, spaced at approximately two full helical turns downstream of USS, are anticipated to contribute to DNA melting and facilitate kinking of the DNA helix in the USS core motif to accommodate DNA entry through the outer membrane pore (14). The DUS-binding protein is the type IV minor pilin ComP (22–24) whereas the USS-specific protein remains unidentified. Since the components of the type IV pilus (T4P) machinery are conserved between Neisseriaceae and the Pasteurellaceae it has been proposed that candidates for USS-specificity may be found at the T4P tip or in the many surface exposed proteins (14). Genome wide sequencing maps of DNA uptake in *H. influenzae* demonstrated that USS bias the DNA uptake step which is initiated by extracellular DNA binding (20). Further support for the general involvement of minor type IV pilins in transformation (specific and nonspecific) competence was found in the γ-Proteobacterium *L. pneumophila* (25). Here, the minor type IV pilin FimT was shown important for DNA uptake in *L. pneumophila* and also DNA binding in *Pseudomonas aeruginosa* and *Xanthomonas campestris* (25). FimT orthologs were also identified across a range of γ-Proteobacteria and particularly common FimT representatives were identified in Xanthomonadales, Alteromonadales and Pseudomonadales (25). Like ComP, the 3D structure FimT displays an electropositive stripe which was found associated with DNA binding (25). The existence of a DNA-binding receptor on the *H. inf* surface was deduced more than half a century ago and this study set out to comprehensively explore USS-receptor candidates using a structural modeling approach involving the latest deep-learning tools.

## Materials and methods

The search for likely USS-receptor candidates was initiated with the download of all available Pasteurellaceae genomes from NCBI in FASTA/GFF3 format (Fig. 1).

**Figure 1.**
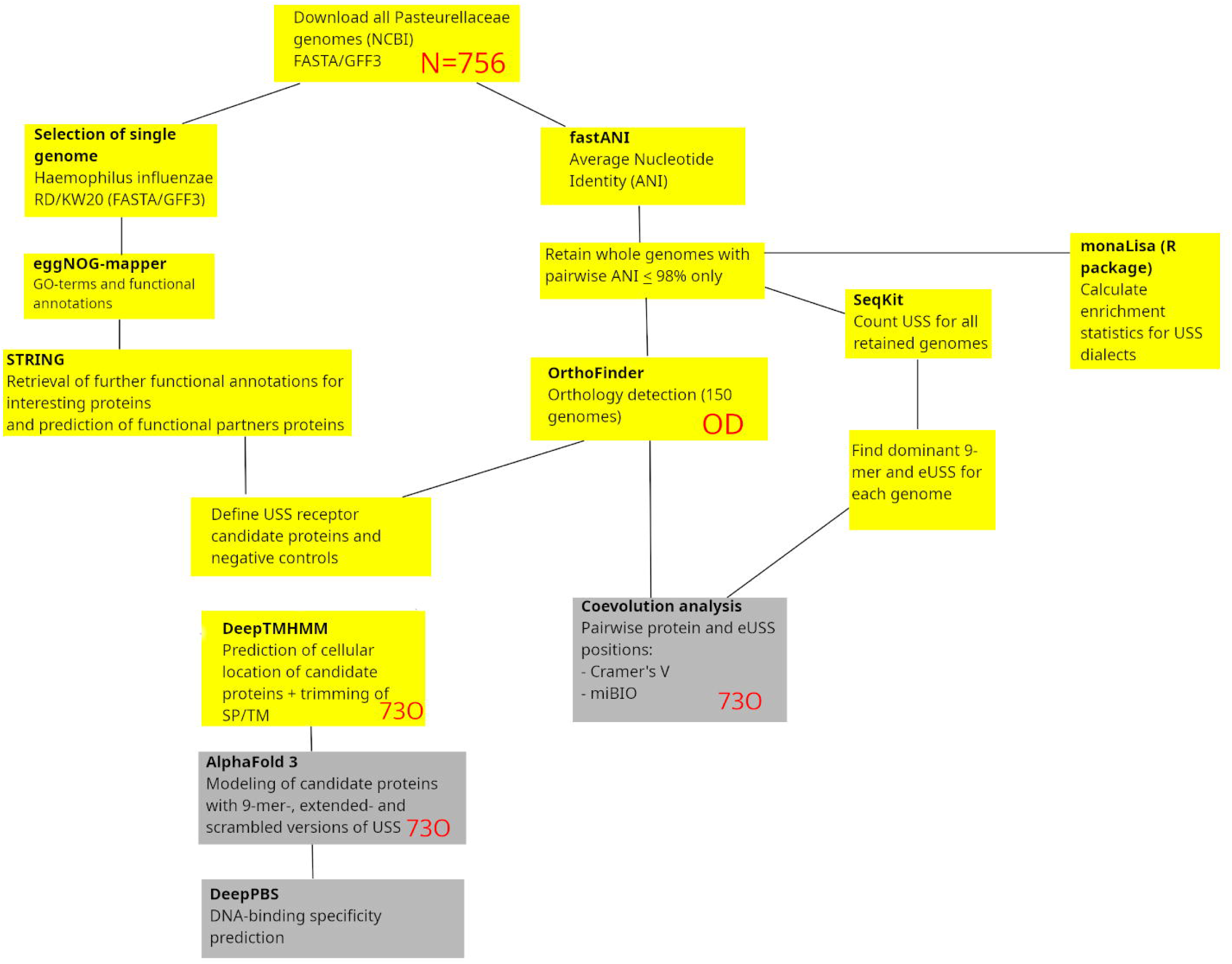
Methodological pipeline in the study. Starting with the download of all Pasteurellaceae genomes, the study takes two connected routes, one leading to coevolution analysis and the other to protein-DNA structural modeling and DNA-binding specificity predictions. Outputs from the yellow boxes are preparatory outputs reported in the Supporting information, whereas the grey boxes are results reported in the Results section. SP = Signal peptides. TM = Transmembrane helices. Red letters refer to analysis output/input.

### eggNOG-mapper and STRING - discriminating unlikely USS receptor candidate proteins

eggNOG-mapper v2.1.12 (26,27) was used to assign all proteins in the *H. influenzae* Rd (H. inf Rd) proteome to orthologous groups in the eggNOG database (Fig. 1). Diamond v. 2.1.10 (28) was employed as the sequence search aligner, set to ultra-sensitive mode to maximize homolog retention. The option to annotate with experimental and non-electronic (--all) Gene Ontology (GO) evidence was used to ensure comprehensive functional classification. To save computational time and also optimize accuracy of annotations, the parameters --tax_scope and --tax_scope_mode were set to Gammaproteobacteria and Bacteria, respectively. The output was then screened and filtered applying GO terms and functional descriptions considered relevant to USS receptor function listed in Table S1.

This annotated eggNOG output contained functional annotations for 296 H inf Rd proteins, which were then queried against the STRING database using an in-house script (https://github.com/stianale/USS-receptor) to retrieve further functional annotations and top-ranked interaction partners (Table S2).

### fastANI and OrthoFinder

To identify orthologs of USS receptor candidate proteins, fastANI v. 1.34 (29) was used to calculate the average nucleotide identity (ANI) across all available (N = 756) *Pasteurellaceae* whole genomes as of Dec 4, 2024, using the many-to-many approach as per instructions in the fastANI GitHub repo (https://github.com/ParBLiSS/FastANI). Using an in-house script, genomes were clustered based on a minimum reciprocal ANI of 98%, and for each cluster one representative genome for each unique annotated species was retained, retaining 150 genomes for downstream analysis. OrthoFinder v. 3.0.1b1 (30) was then run on this dataset using Diamond v. 2.1.10 (28) as the sequence search engine and the option to infer gene trees from MAFFT v. 7.525 (31,32) sequence alignments, referred to as the OrthoFinder dataset (OD) was used. NCBI genome accessions for the OD genomes are listed in Table S2. USS receptor candidacy was then assessed and filtered based on (i) functional relevance in eggNOG and STRING annotations, and (ii) orthogroup coverage (number of OD genomes with ortholog present) of at least 90% in the OrthoFinder dataset (OD). After applying these filters, the final number of USS receptor candidates was reduced to 15 proteins (15P) as highlighted in Table S2, while 281 of the initial eggNOG orthogroups were deemed unlikely to be the USS receptor of which a selected set were used as negative controls in modeling (see below).

### DeepTMHMM, AF3 and DeepPBS

DeepTMHMM is a deep-learning protein language model-based tool which predicts the topology of transmembrane proteins (31) and provides information which can educate understanding of cellular localization of proteins. DeepTMHMM v. 1.0.24 was used to predict the cellular localization of orthologs from 10 selected OD species for the USS candidates and negative controls listed in Table S2, five species each highly enriched in Hin-USS or Apl-USS. Since USS-binding takes place prior to passage of the outer membrane into a DNase-protected stage (20) and current natural transformation models show extracellular DNA binding (8), we were interested in domains predicted to face the exterior environment. The DeepTMHMM predictions were thus used to guide modeling of orthologous protein domains with DNA in AF3. When DeepTMHMM predicted a single extracellular domain for an ortholog, this domain was selected for AF3 modeling. In cases where an ortholog was predicted with multiple extracellular domains, or the extracellular domain in single-domain proteins was shorter than 33 residues, well below the limit of known functional domains (33), the full length protein sequences were used for AF3 modeling.

The orthologous proteins and domains were then modeled in AF3 together with the species’ dominant 9-mer USS (orthogroup_USS_) in addition to scrambled forms of the USS (orthogroup_scr_), each running a minimum of 20 replicates with different seeds. The reasons for doing this were two-fold: 1. Discriminate on modeled USS-binding specificity - expecting statistically significant relative higher confidence metrics for orthogroup_USS_ predictions than orthogroup_scr_ predictions for the true USS receptor than other orthogroups. 2. Discriminate on modeling robustness - expecting higher confidence metrics for the true USS receptor relative to other orthogroups. For the AF3 output models, an interface predicted template modeling (ipTM) range of > 0.6 <= 1 (ipTM_r_) was used to determine potential DNA binding properties of proteins, following . Furthermore, we ran ipTM-, PAE-, CPPM and pLDDT-wise Wilcoxon rank sum tests on orthogroup_USS_ vs. orthogroup_scr_ results for predictions within ipTM_r_ as previously described (5). Based on strength of interactions between the proteins and the DNA as measured by the lower cut-off ipTM value of 0.6 (34), as well as a lower predicted aligned error (PAE) for orthogroup_USS_ compared to orthogroup_scr_, we identified the strongest USS-specific binding orthogroup candidate to be prepilin peptidase dependent protein A (PpdA). Different names assigned PpdA in *H. inf* Rd. were Uncharacterized protein HI0938 (Swiss-Prot), pilus assembly FimT family protein (RefSeq), prepilin-type cleavage/methylation domain-containing protein (RefSeq), type II secretion system protein (INSDC), type II secretion system GspH family protein (INSDC) and Tfp pilus assembly protein FimT/FimU (INSDC). The HI0938 gene had previously been given the name *comN* (35).

DockQ (36) was calculated for pairwise orthogroup_USS_ top models (ipTM) for all 10 modeled Pasteurellaceae species before a graph-edge method was used to place models in clusters where all members have reciprocal DockQ >= 0.490. Using 0.490 as a lower DockQ threshold instead of 0.8 was a good compromise for cluster size and number, which we wanted to increase and decrease, respectively. No models were permitted to be allocated in multiple clusters. For each species, the models were superimposed in the largest clusters to the α-carbons and visualized in overlays using PyMOL v. 3.0.0 Open Source (37) (Figure S1).

We explored if the geometric deep-learning algorithm, DeepPBS v. 1.0 (6), could predict the DNA binding specificity of the AF3 predicted USS receptor to match the USS motif across distinct Pasteurellaceae family orthologs. From the protein structure DeepPBS aggregates the atom environment (type, charge, radius) which is laid upon a symmetrized DNA helix structure (sym-helix) to which a bipartite geometric convolution is made. This overall convolution is informed by DNA groove and DNA shape readouts consisting of major and minor groove convolutions (Groove readout, ie. not base readout) and DNA backbone sugar and phosphate convolutions (Shape readout). We ran DeepPBS using the default module which considers shape and groove properties of the DNA (“both readout”). We reasoned that if this approach predicted binding specificity to match the two different USSs (Hin-USS and Apl-USS), then this would be additional support for the AF3 modeling as informative on the Pasteurellaceae USS receptor protein. Assuming that the DNA-binding specificity of the USS-receptor would be reflected as an evolutionary trace in the genomic conservation of each nucleotide in the USS, DeepPBS predictions and sequence logos of genomic USS conservation were compared. The sequence logos were made as devised in (21) allowing 1 mismatch to the 9-mer USS. Finding correlations between USS conservation and DeepPBS predicted specificity could further establish a functional connection between predicted specificity and the existing genomic USS imprint consensus. Finally, in a series of negative control experiments we explored if DeepPBS could predict any DNA-position relative nucleotide preference *de novo* using modeled PpdA-DNA complexes with scrambled USS-sequences to suggest nucleotide biases independent of the USS.

DeepPBS outputs probability distributions of the four nucleotides in each position of one DNA strand (Watson) of the input protein-DNA complex, heavy-atom relative importance scores for the protein and a nucleotide binding-specificity plot for an input protein-DNA complex. The mean predicted nucleotide probabilities at each USS position from ten USS-receptor candidates were calculated separately and the outputs were combined according to USS-dialect (Hin-USS and Apl-USS). Statistical significance was tested using a one-sample t-test to check for mean predicted nucleotide probabilities significantly deviating from 0.25 (random chance for any nucleotide) at each USS position. Ensembles of DeepPBS nucleotide binding-specificity plots were animated by first converting SVG to PNG using inkscape v. 1.4-dev (38) and then concatenating PNGs to GIF format using ImageMagick v. 6.9.12-98 (39) (Video S1 and Video S2).

### USS statistics, extended USS and coevolution analysis

To inform the structural modeling, prediction of sequence-specificity and coevolution analyses, the per-genome numbers of Hin-USS and Apl-USS of all OD genomes were extracted using SeqKit v. 2.3.0 (33) and corrected for genome size (counts/Mbp) (Table S3). For the coevolution analysis specifically, we widened the USS to 17 nt extending into up- and downstream less conserved, yet potentially evolutionarily informative regions. Individual nt positions in the eUSS were numbered 1-17. The genome-specific extended USS (eUSS) were calculated from an alignment of all genomic occurrences of the dominant 9-mer USS (Hin-USS/Apl-USS) including one upstream and seven downstream additional nts. Regarding specific orthogroup-eUSS pairwise positions, only the eUSS positions in the alignment that were variable had potential for detecting coevolved pairs in the coevolution analyses (see below). eUSS positions 2, 5-9 and 15-16 are invariable and hence not informative.

Pasteurellaceae genomes statistically overrepresented in either USS dialect (Hin or Apl) were considered informative for the orthogroup-USS coevolution analysis. Thus, enrichment statistics for the USS dialects were calculated for all OD genomes (Table S3). A Markov model of order 4 was used, following the recommendations in Davidsen et al. (40), although we used 9-mers instead of 10-mers. All OD genomes were found to be overrepresented by either Hin-USS or Apl-USS to variable extent (from 31.34/mb to 913.03/mb shown in Table S3), and were included in the coevolution analysis. Multiple sequence alignments (MSA) for the 73 AF3 modeled orthogroups (73O) were compared to MSAs of the calculated eUSS (Watson strand) in a systematic manner, using two different approaches.

(#1) Cramér’s Φ was calculated for pairwise sites across orthogroup alignments and eUSS. Bonferroni correction of p-values was applied to account for multiple testing. Only the OD genomes having orthologs in each respective orthogroup were included in the calculation of Φ*_c_*. For orthogroups containing multiple copies for a species (paralogs), underscores were appended to the genome accession in both the orthogroup and eUSS MSAs to distinguish paralog 1, 2, etc. Φ*_c_* was calculated for all pairwise positions in the two alignments as a measure of coevolution between the orthogroups and eUSS.

(#2) miBio (41,42) was employed to calculate mutual information (MI) values across pairwise positions across orthogroup alignments and eUSS. The option to shuffle the identities of the residues and subtract shuffled MI values from original ones was applied. As miBio allows for customization of grouping of residues, we used a custom grouping pairing amino acids D/E, K/R and N/Q, because these have been shown to have comparable nucleotide-binding preferences in protein-DNA complexes (43,44), as well as treating gaps as separate, valid states. Finally, the mean MI for the resulting positive MI values for each orthogroup was calculated.

## Results

The search for the USS-receptor went through a series of initial steps as described in the Material and methods section and the outputs are reported in the Supporting data (Table S1-S3; Presentation S1-2), involving functional gene ontology assessment, cellular location predictions, USS-enrichment statistics and orthogroup coverage across USS-enriched genomes/species. These outputs were used in consecutive deep-learning structural modeling of protein-DNA complexes (refer to Table S2 for all orthogroups modeled in AF3), DNA-binding specificity predictions and coevolutionary analysis described in each section below.

### Assessment of Protein-DNA complexes modeled in AF3

Out of 73 modeled orthogroups, AF3 modeled only six potential candidate DNA-binding orthogroups across all ten species with at least one model within ipTM_r_. These were: PpdA, DUF4198 domain-containing protein, two hypothetical proteins, protein annotated “MULTISPECIES: hypothetical protein”, VirK/YbjX family protein and YchJ family protein. Fig. 2 shows the ipTM distribution of all AF3 models generated within ipTM_r_ for all six orthogroups modeled with either USS (orthogroup_USS_) or scrambled versions of USS (orthogroup_scr_). The rationale for comparing the USS and scrambled models was the expectation that AF3 would generate better quality models with USS than scrambled USS for the USS receptor. The statistical comparisons are reported in CodeOutput S1. The distributions of the three other modeling quality parameters pLDDT, PAE and CPPM are plotted in Fig S1-3. Since all ten modelled species had USS enriched genomes and expected to have the USS receptor, the OrthoFinder orthogroup coverage (Table S2) for all six orthogroups were recorded these and in the 150 OD genomes (Table S3) as described in each section below. The AF3 modeling results of the six orthogroups were as follows:

**Figure 2.**
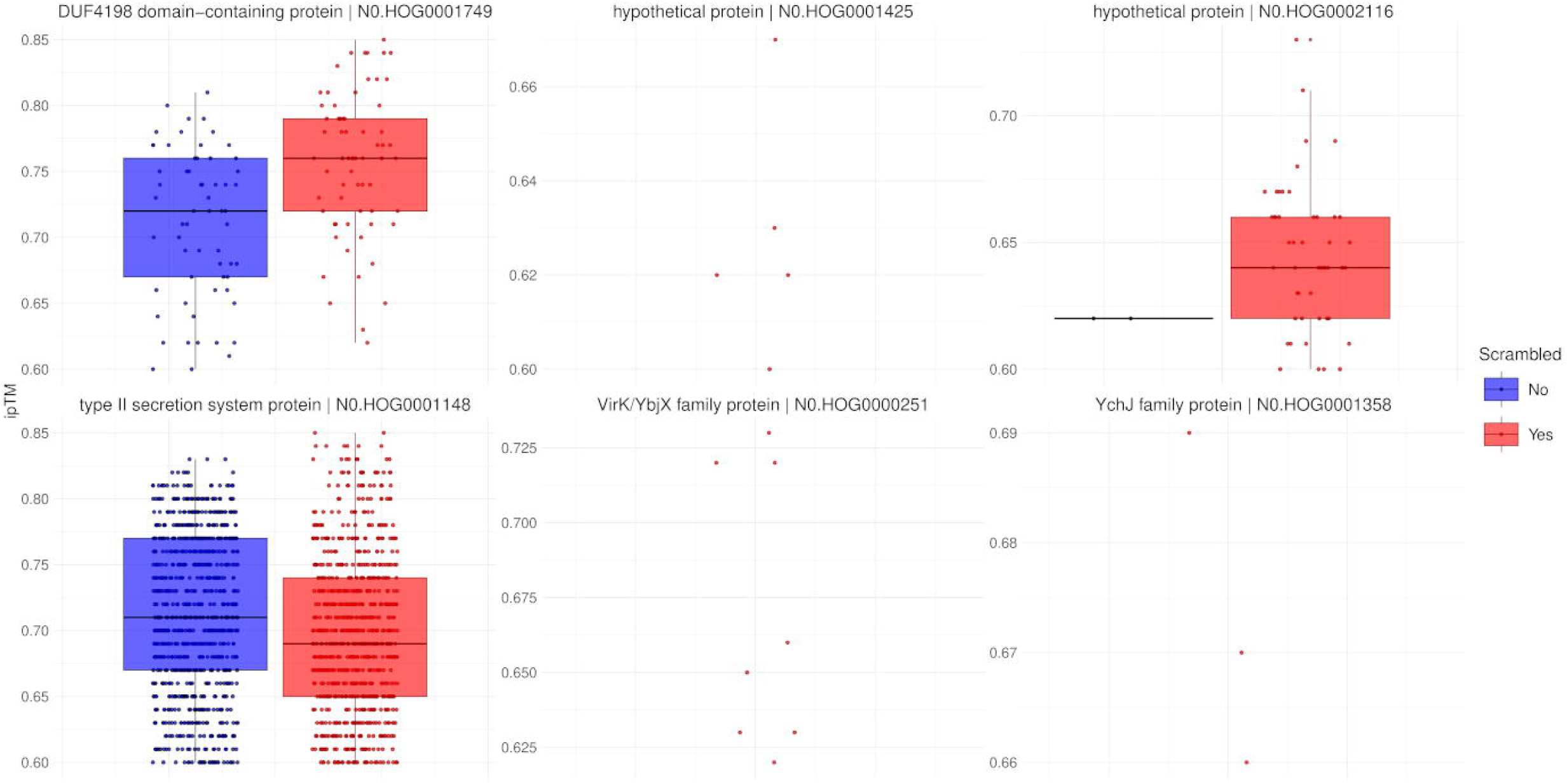
Distribution of model confidence ipTM values. All AF3 orthogroup_USS_ and orthogroup_scr_ models for the six orthogroups with results within ipTM_r_ represented as box plots.

#### PpdA (type II secretion system protein)

1629/2000 (81.45 %) models fell within ipTM_r_, (Fig. 2) and the orthogroup coverage was 10/10 investigated species and 147/150 OD species.

10/10 investigated species had models in ipTM_r_. ipTM and pLDDT were significantly higher for PpdA_USS_ than PpdA_scr_ (Wilcoxon rank sum test, p-value = 2.43×10−7; p-value = 6.36×10−5). PAE was not significantly different between PpdA_USS_ and PpdA_scr_ (Wilcoxon rank sum test, p-value = 0.808). CPPM was significantly lower for PpdA_USS_ than PpdA_scr_ (Wilcoxon rank sum test, p-value = 0.0107). Structural overlays of the top-ranking PpdA_USS_ models in the ten modeled species are shown in Fig. S5 and structural overlays of the largest cluster of *H. inf* Rd PpdA_Hin-USS_ AF3 models in Fig. S6.

#### Hypothetical protein (OrthoFinder orthogroup N0.HOG0002116)

51/400 (12.75%) models were within ipTM_r_ (Fig. 2), and the orthogroup coverage was 2/10 investigated species and 44/150 OD species. 2/10 species had models within ipTM_r_. ipTM and pLDDT were non-significantly and significantly higher for N0.HOG0002116_scr_ than N0.HOG0002116_USS_, respectively (Wilcoxon rank sum test, p-value = 0.178; p-value =0.0392), while PAE and CPPM were significantly and non-significantly lower for N0.HOG0002116_scr_ than N0.HOG0002116_USS_, respectively (Wilcoxon rank sum test, p-value = 0.0392; p-value = 0.19) (Fig S1-3; Code Output S1).

#### DUF4198 domain-containing protein

128/1600 (8.0 %) models were within ipTM_r_, (Fig. 2) and the orthogroup coverage was 8/10 investigated species and 77/150 OD species. 2/10 species had models within ipTM_r_. The DUF4198_scr_ models had significantly higher ipTM than DUF4198_USS_ models (Wilcoxon rank sum test, p-value = 1.56×10−5). PAE and CPPM were significantly lower for DUF4198_scr_ than DUF4198_USS_ (Wilcoxon rank sum test, p-value = 2.26×10−4; p-value = 1.19×10−4). pLDDT was significantly higher for DUF4198_scr_ than DUF4198_USS_ (Wilcoxon rank sum test, p-value = 5.51×10−8) (Fig 3 A-D; Code Output S1).

**Figure 3.**
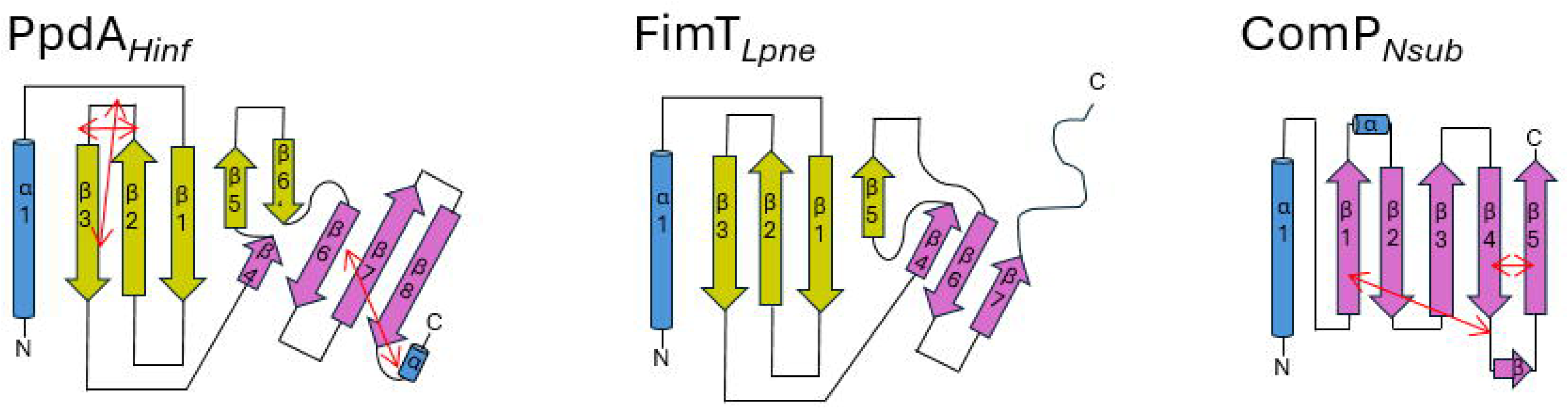
Topology diagrams of PpdA, FimT and ComP. The common α-helix (truncated) at the N-terminal of the proteins is colored in blue, and β-sheets in yellow and purple. Short α-turns and β-strands are indicated. Disulfide bridges are shown with double red arrows. The diagrams and coloring are based on the FimT*_Lpne_* described in Braus et al. (25)

#### Hypothetical protein (OrthoFinder orthogroup N0.HOG0001425)

5/1800 (0.28 %) models fell within ipTM_r_ (Fig. 2) and the protein had an orthogroup coverage of 9/10 investigated species and 120/150 OD species. All models within ipTM_r_ were orthogroup_scr_. 2/10 species had models in ipTM_r_ (Fig S1-3; Code Output S1).

#### VirK/YbjX family protein

8/1600 (0.5%) AF3 models were within ipTM_r_ (Fig. 2), all being VirK/YbjX_scr_ models, being represented by 4/10 investigated species. The orthogroup coverage was 9/10 modeled species and 142/150 OD species. All models within ipTM_r_ were orthogroup_scr_. (Fig S1-3; Code Output S1).

#### YchJ family protein

3/2000 (0.15%) models were within ipTM_r_ (Fig. 2), even though this protein had a strong orthogroup coverage of 10/10 investigated species and 140/150 OD species. 2/10 species had models in ipTM_r_. All models within ipTM_r_ were orthogroup_scr_. (Fig S1-3; Code Output S1).

Having found the robust modeling support for PpdA as the USS-receptor described above, we explored further the structural similarities between PpdA and the DUS-receptor ComP in Neisseriaceae (5,24,45,46) and the non-specific DNA-receptor in *L. pneumophila* (25) (Fig 3 and Fig S5). The three proteins PpdA*_Hinf_*, FimT*_Lpne_* and ComP*_Nsub_* share the characteristic N-terminal α-helix and a large globular domain built over β-sheet(s). PpdA*_Hinf_* shares the overall structural fold with FimT*_Lpne_*. The globular domains of PpdA*_Hinf_* and FimT*_Lpne_* are considerably larger than ComP*_Nsub_*, consisting of 7-8 β-strands in two differently angled β-sheets as opposed to ComP’s singular β-sheet. The *H.inf* Rd PpdA differs from FimT*_Lpne_* with one short β-strand (β-6’) and one additional long beta-strand β8 at the C-terminus. PpdA*_Hinf_* has three disulfide bridges (Cys-Cys) whereas FimT*_Lpne_* has none. ComP*_Nsub_* differs from PpdA*_Hinf_* by having a single β-sheet of five strands interconnected with two disulfide bridges. The two disulfide bridges in ComP have been shown essential for DUS-specific binding (24) and one of these connects a long loop (DD-loop) across the β-sheet onto β1 (Fig. 3). PpdA*_Hinf_* similarly connects its C-terminal loop across the β-sheet to β6. In the AF3 top-scoring PpdA_Hin-USS_ models and other PpdA_USS_ models with ipTM ≥ 0.6/DockQ ≥ 0.49, this loop is modeled to interact with DNA (Figs. S4 and S6). One typical overall DNA binding mode was found in top scoring Ppda_USS_ models of both Hin-USS and Apl-USS species (Fig. S4).

### DeepPBS DNA-binding specificity predictions for Hin-USS species

DeepPBS was run on 808 PpdA_USS_, 797 PpdA_scr_ AF3 models which were within ipTM_r_. We present the DeepPBS results using the default module (“both readout”). The DeepPBS results for each species were as follows:

Predictions of PpdA_Hin-USS_ models of *H. inf* Rd modeled with variable significance (* p<0.05, ** p<0.01 and *** p<0.001) overrepresented nucleotides matching Hin-USS in positions 1(A), 2(A), 3(G), 4(T), 5(G), 6(C), 7(G) and 8(G) (Fig 4 A; Table S8). Position 9(T) was non-significantly different from random, yet on the overrepresented side together with the significantly (***) overrepresented pyr-pyr transition 9(C). Both permutational transversions 9(G) and 9(A) were significantly (***) underrepresented. The descending order of mean predicted probabilities for each Hin-USS nucleotide was 4(T), 6(C), 8(G), 2(A), 7(G), 1(A), 5(G), 3(G) and 9(T). The sequence logo of Hin-USS from this species showed that the weakest predicted 9(T) also to be the least conserved nucleotide relative to the other USS positions and the significantly (***) underrepresented 9(G) to be the rarest permutation (Fig. 4 A)

**Figure 4.**
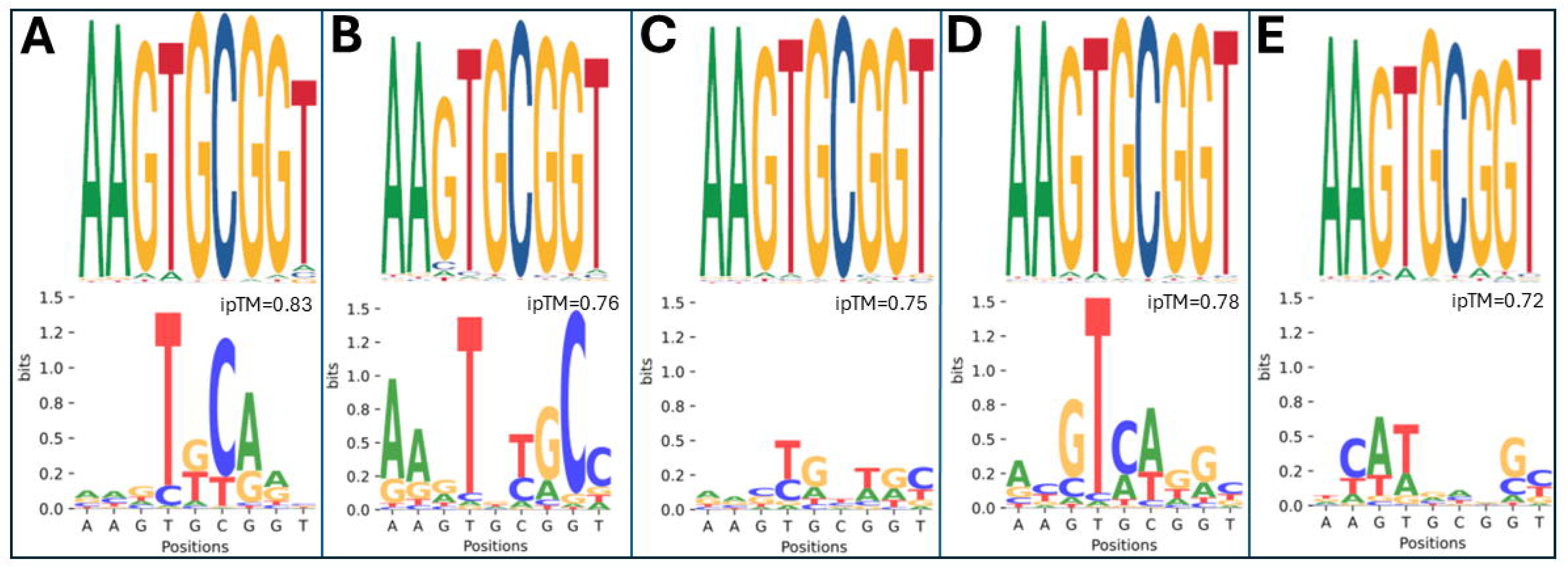
Hin-USS conservation and DeepPBS predictions. Sequence logos of genome specific Hin-USS conservation (top panels) and mean predicted nucleotide frequencies in bits score from DeepPBS predictions on top-scoring (max ipTM) PpdA_Hin-USS_ AF3 inputs (lower panels). ipTM values for each of the AF3 input models are shown. **A:** *H. influenzae* Rd. **B**. *Mannheimia succiniciproducens* strain MBEL55E. **C**: *Aggregatibacter actinomycetemcomitans* strain 31S. **D**: *Aggregatibacter* sp oral taxon 513. **E**: *Pasteurella multocida* strain NCTC8282.

AF3 models of PpdA_Hin-USS_ models of *Mannheimia succiniciproducens* strain MBEL55E predicted with variable significance (* to ***) overrepresented nucleotides matching Hin-USS in all positions 1-9 (Fig 4 B; Table S8). Positions 3(G) and 5(G) had weaker overrepresented significance (* and **) than the other six nucleotide positions (***). The descending order of mean predicted probabilities for each Hin-USS nucleotide was 1(A), 4(T), 6(C), 9(T), 2(A), 7(G), 8(G), 5(G) and 3(G). The sequence logo of the USS for this species showed that the weakest predicted position 3(G) to also be markedly less conserved relative to all other Hin-USS positions and notably relative to the other Hin-USS species. (Fig. 4 B)

AF3 models of PpdA_Hin-USS_ of *A. actinomycetemcomitans* strain 31S predicted with strong significance (***) overrepresented nucleotides matching Hin-USS in position 1(A), 2(A), 4(T) and 8(G) (Fig 4 C; Table S8). In contrast, position 3(G) to match Hin-USS was significantly (**) underrepresented and the complementary pur-pyr permutational transversion 3(C) was significantly (***) overrepresented. Positions 5(G), 6(C) and 7(G) were non-significantly different from random and the complementary permutational transversions 5(C), 6(G) and 7(C) were significantly (** and ***) underrepresented. The descending order of mean predicted probabilities for each USS nucleotide was 4(T), 8(G), 1(A), 2(A), 5(G), 7(G), 6(C), 9(T) and 3(G). The sequence logo of the Hin-USS in this species showed that the weakest and predicted underrepresented position 3(G) was also among the least conserved of the Hin-USS positions (Fig. 4 C).

AF3 models of of PpdA_Hin-USS_ of *Aggregatibacter sp.* oral taxon 513 predicted with variable significance (* to ***) overrepresented nucleotides matching Hin-USS in positions 1(A), 2(A), 3(G), 4(T), 7(G), 8(G) and 9(T) (Fig 4 D; Table S8). Positions 5(G) and 6(C) were non-significantly different from random, yet on the overrepresented side together with their significantly overrepresented permutational transitions 5(A)* and 6(T)***, respectively. The descending order of mean predicted probabilities for each Hin-USS nucleotide was 4(T), 7(G), 8(G), 1(A), 3(G), 2(A), 9(T), 6(C) and 5(G). Although non-significantly predicted by DeepPBS, positions 5(G) and 6(C) are among the best conserved nucleotides relative to the other USS positions in the USS sequence logo in this species (Fig. 4 D).

AF3 models of PpdA_Hin-USS_ models of *P. multocida* strain NCTC8282 predicted with strong significance (***) overrepresented nucleotides matching Hin-USS in positions 1(A), 3(G), 4(T) and 9(T) (Fig 4 E; Table S8). All other positions 2(A), 5(G), 6(C), 7(G) and 8(G) were non-significantly different from random. The descending order of mean predicted probabilities for each Hin-USS position was 4(T), 3(G), 9(T), 1(A), 6(C), 2(A), 5(G), 8(G) and 7(G). In the sequence logo, the weakest predicted position 7(G) was also the least conserved relative to the other Hin-USS positions and in this species also relative to the other Hin-USS species. Also, the *P. multocida* NCTC8282 USS differed from the other Hin-USS sequence logos by having a relatively less conserved 6(C) and a particularly conserved 9(T) which was uniquely better conserved than 7(G) and 8(G) in this species. It is notable that the weakest predicted and least conserved position in *H. inf* Rd 9(T) was both predicted with strong significance and was better conserved in *P. multocida* (Fig. 4 E).

Considering similarities in the predictions for each species (Fig 4 A-E, *H. inf* Rd PpdA_Hin-USS_ and *A. sp.* oral taxon 513 PpdA_Hin-USS_ were most similarly predicted across Hin-USS with particularly strong predictions for 4(T), 6(C), 7(G) and 8(G), also similar to *M. succiniciproducens* strain MBEL55E with a strong additional 1(A) prediction. When considering the average predictions of nucleotide representation from all five Hin-USS species, the overrepresented nucleotides matching Hin-USS was found with strong significance (***) above random (25%) in all nine positions (Fig 5; Table S8; Video S1). The nucleotide probabilities for each USS position ranged from highest 0.6448 for 4(T) to lowest 0.2917 for 3(G). Position 1(A), 2(A) and 4(T) were predicted without any significant ambiguities, whereas all other positions were predicted together with one other significant ambiguity, transversion permutations in positions 3(G/C) and 5(G/T) and transition permutations in positions 6(C/T), 7(G/A), 8(G/A) and 9(T/C). In descending order, the highest levels of overrepresentation were found in positions 4(T), 1(A), 6(C), (8G), 7(G), 2(A), 9(T), 5(G) and 3(G). DeepPBS runs with the Hin-USS orthogroup_scr_ models show that all predictions failed to predict *de novo* a coherent signal different from random (Table S9).

**Figure 5.**
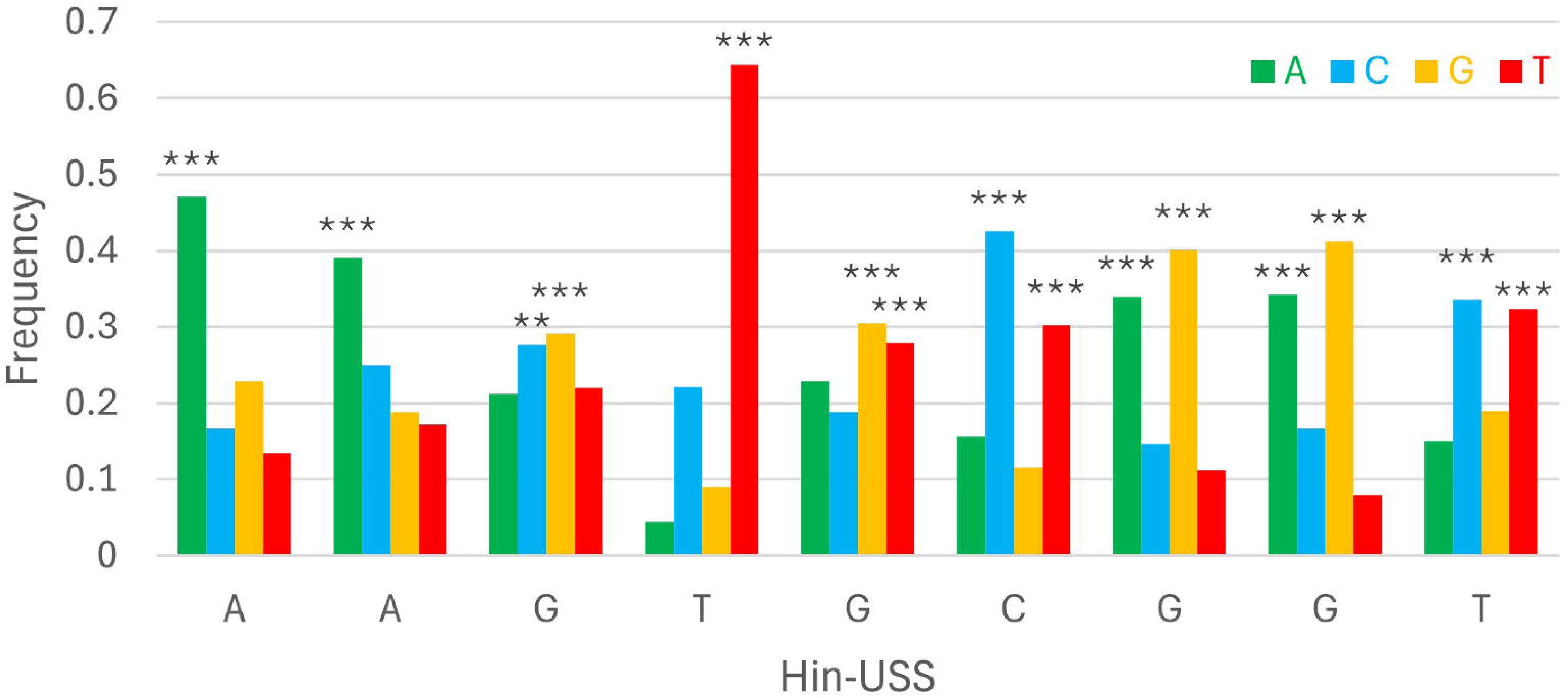
Consensus DeepPBS predictions of PpdA specificity against the reference Hin-USS. Distribution of consensus predicted nucleotide probabilities in each of the nine Hin-USS positions. Asterisks denote significant representation above random (0.25), ***P<0.001, **P<0.01, *P<0.05.

### DeepPBS DNA-binding specificity predictions for Apl-USS species

AF3 models of PpdA_Apl-USS_ of *Actinobacillus equuli* subsp. haemolyticus strain 3524 predicted with strong significance (***) overrepresented nucleotides matching Apl-USS in positions 1(A), 2(C), 3(A), 4(A), 6(C), 7(G), 8(G) (Fig 6 A, Table S8). Position 5(G) to match Apl-USS was underrepresented with weaker significance (*) below random chance (25%) and the non-complementary transversion permutation 5(T) was overrepresented with strong significance (***). Position 9(T) to match Apl-USS was underrepresented with strong significance (***) and the non-complementary transversion permutation 8(G) was overrepresented (***). The descending order of mean predicted probabilities for each Apl-USS nucleotide was 1(A), 3(A), 7(G), 8(G), 2(C), 4(A), 6(C), 5(G) and 9(T). The USS sequence logo from this species showed that the significantly overrepresented positions 3(A), 4(A), 6(C), 7(G) were also the most conserved nucleotides relative to the other USS positions. In contrast, positions 1(A) and 2(C) were the least conserved, yet significantly overrepresented (***). Underrepresented position 5(G) was less conserved than the adjacent 3(A), 4(A), 6(C) and 7(G) pairs relative to the other Apl-USS species except *Actinobacillus lignieresii* strain NCTC4189 with a resembling sequence logo (Fig 6 A-E).

**Figure 6.**
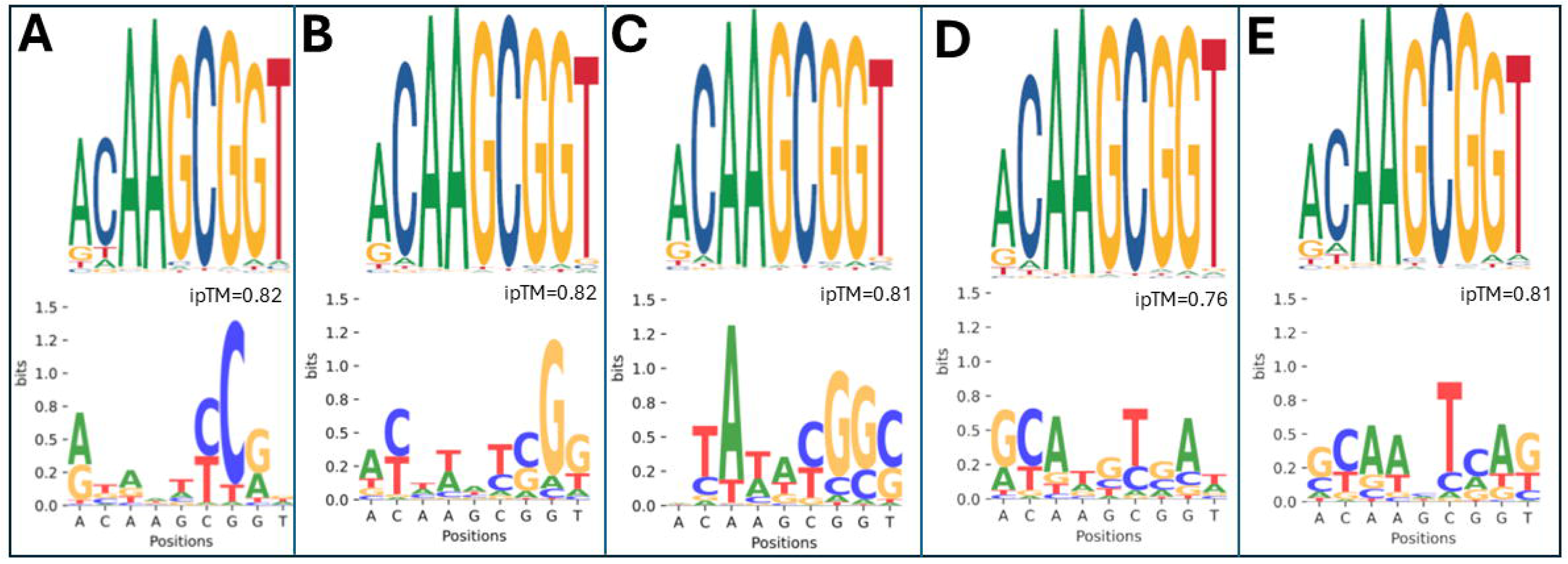
Apl-USS conservation and DeepPBS predictions. Sequence logos of genome specific Apl-USS conservation (top panels) and mean predicted nucleotide frequencies in bits score from DeepPBS predictions on top-scoring (max ipTM) PpdA_Apl-USS_ AF3 inputs (lower panels). ipTM values for each of the AF3 input models are shown. **A**: *Actinobacillus equuli subsp.* haemolyticus strain 3524. **B**: *Frederiksenia canicola* strain HPA 21. **C**: Pasteurellaceae bacterium Orientalotternb1. **D**: *Mannheimia bovis* strain 39324S-11. **E**: *Actinobacillus lignieresii* strain NCTC4189.

AF3 models of PpdA_Apl-USS_ of *Frederiksenia canicola* strain HPA 21 predicted with strong significance (***) overrepresented nucleotides matching Apl-USS in positions 1(A), 2(C), 3(A), 4(A), 6(C), 7(G) and 8(G) (Fig 6 B, Table S8). Positions 5(G) and 9(T) were underrepresented with strong significance (***). In position 5(G) both the complementary transversion permutation 5(C) and the transition permutation 5(A) were significantly (***) overrepresented. In position 9(T) the non-complementary transversion permutation 9(G) was significantly (***) overrepresented. The descending order of mean predicted probabilities for each Apl-USS nucleotide was 8(G), 7(G), 1(A), 2(C), 4(A), 3(A), 6(C), 9(T) and 5(G). The USS sequence logo from this species showed that the significantly overrepresented positions 3(A), 4(A), 6(C), 7(G), 8(G) were also the most conserved nucleotides relative to the other USS positions and together with the conserved yet underrepresented 5(G). 1(A), 2(C) and 9(T) were less conserved than the above and 1(A) the least (Fig 6 A-E).

AF3 models of PpdA_Apl-USS_ models of Pasteurellaceae bacterium Orientalotternb1 predicted with strong significance (***) overrepresented nucleotides matching Apl-USS in positions 2(C), 3(A), 4(A), 7(G) and 8(G) (Fig 6 C, Table S8). Position 1(A) was underrepresented with weak significance (*) and the position significantly overrepresented with the non-complementary transversion permutation 1(C)* and the transition permutation 1(G)***.

Position 6(C) was underrepresented with strong significance (***) and the position significantly overrepresented with the transition permutation 6(T)***. Positions 5(G) and 9(T) were non-significantly different from random. The descending order of mean predicted probabilities for each Apl-USS nucleotide was 2(C), 3(A), 4(A), 8(G), 7(G), 6(C), 9(T), 5(G) and 1(A). The USS sequence logo from this species showed that the weakest predicted and underrepresented 1(A) was also the least conserved. In contrast, the strongest predicted 2(C) was less conserved than positions 3-9 (Fig 6 A-E).

AF3 models of PpdA_Apl-USS_ models of *Mannheimia bovis* strain 39324S-11 predicted with strong significance (***) overrepresented nucleotides matching Apl-USS in positions 2(C), 3(A), 4(A), 7(G) and 9(T) (Fig 6 D, Table S8). Positions 1(A) and 5(G) were underrepresented with strong significance (***). In position 1(A) the transition permutation 1(G) was significantly (***) overrepresented. In positions 5(G) both the complementary transversion permutation 6(T) and the transition permutation 6(A) were significantly (**) overrepresented. 6(C) and 8(G) were non-significantly different from random and both positions significantly overrepresented with their respective transition permutations 6(T) and 8(A), respectively. The descending order of mean predicted probabilities for each Apl-USS nucleotide was 2(C), 3(A), 4(A), 9(T), 7(G), 6(C), 8(G), 5(G) and 1(A). The USS sequence logo from this species showed that both the weakest predicted 1(A) and the strongest predicted 2(C) were less conserved than positions 3-8 and with 1(A) the least. A unique feature of this species’ sequence logo was that position 7(G) and 9(T) were equally conserved where the other Apl-USS sequence logos showed better conservation for 7(G) than 9(T) (Fig 6 A-E).

AF3 models of PpdA_Apl-USSS_ models of *Actinobacillus lignieresii* strain NCTC4189 predicted with strong significance (***) overrepresented nucleotides matching Apl-USS in positions 1(A), 2(C), 3(A), 4(A), 6(C) and 9(T) and less significantly positions 7(G)** and 8(G)* (Fig 6 E, Table S8). Position 5(G) to match Apl-USS was non-significantly different from random and no other nucleotide was overrepresented in this position. The descending order of mean predicted probabilities for each Apl-USS nucleotide was 3(A), 2(C), 6(C), 4(A), 1(A), 7(G), 9(T), 8(G) and 5(G). The USS sequence logo from this species resembled that of *A. equuli* subsp. haemolyticus strain 3524 with a less conserved 5(G) relative to the 3(A), 4(A), 6(C) and 7(G), equally conserved 8(G) and 9(T) and less conserved 2(C) relative to the other Apl-USS sequence logos. The least conserved nucleotide position was also in this species 1(A) with a strong prediction (Fig 6 A-E).

When considering the Apl-USS species together, DeepPBS predicted the near exact overrepresented nucleotides matching Apl-USS with variable significance (Fig 7; Table S8; Video S2). Positions 1(A), 2(C), 3(A), 4(A), 6(C), 7(G) and 8(G) were predicted overrepresented with strong significance (***). Notably, positions 2-4 distinguish Apl-USS from Hin-USS and 2(C) was strongly predicted whilst having generally lower levels of conservation across Apl-USS species. Position 9(T) was predicted overrepresented with weak significance (*) together with the significantly (***) overrepresented non-complementing transversion permutation 9(G). Position 5(G) was found significantly underrepresented (***) and the non-complementary transversion permutation 5(T) significantly (***) overrepresented. The descending order of mean predicted probabilities was 2(C), 3(A), 4(A), 6(G), 7(G), 1(A), 6(C), 9(T) and 5(G). The only predicted underrepresented position 5(G) was found relatively less conserved in the USS sequence logos of two species (*A. equuli* subsp. haemolyticus strain 3524 and *A. lignieresii* strain NCTC4189). Although 1(A) was the consistently least conserved it was predicted significantly overrepresented (***) in the PpdA_Apl-USS_. In considering the ambiguity of the two highest mean predicted probability nucleotides of each Apl-USS position, position 1, 2, 3, 6, and 8 were all transition permutations (Pyr-Pyr and Pur-Pur), whereas position 4(A) was alternatively predicted with the complementary transversion 4(T) and position 9(T) with the non-complementary transversion permutation 9(G). Again, DeepPBS runs with the Apl-USS orthogroup_scr_ models show that all predictions failed to predict *de novo* a coherent signal different from random (Table S9).

**Figure 7.**
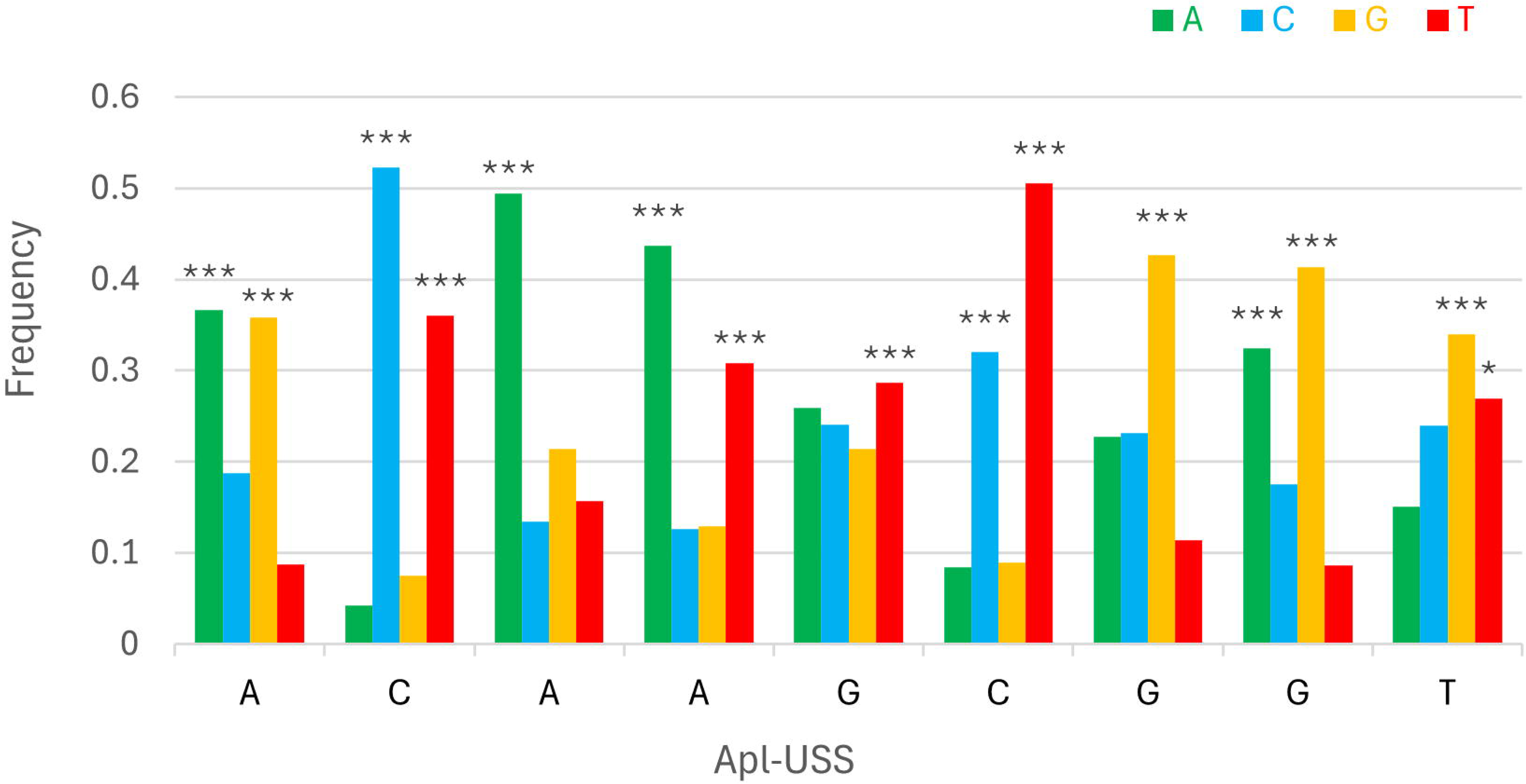
Consensus DeepPBS predictions of PpdA specificity against the reference Apl-USS. Distribution of consensus predicted nucleotide probabilities in each of the nine Apl-USS positions. Asterisks denote significant representation above random (0.25), ***P<0.001, **P<0.01, *P<0.05.

### Orthogroup-eUSS coevolution analyses by Cramér’s Φ and miBIO

To further explore the PpdA candidacy as the USS-receptor we looked for traces of reciprocal influence between orthogroups and eUSS in coevolution analyses. In contrast to modeling, the USS (9-mer) was extended to eUSS (17-mer) as described in the Materials and methods section in order to widen the evolutionary signal with variable DNA positions. Φ*_c_* was calculated for all pairwise orthogroup-eUSS positions For example the PpdA orthogroup alignment (Supplementary Data 1) paired with the informative positions (1, 3-5, 11-14 and 17) in the 17-mer USS provided 996 unique pairs (Table S4-5), which in turn could be assessed for correlation. Of all orthogroups tested, PpdA had the highest Φ*c* factor of 5.05 (Figure 8; Table S4), meaning the number of significantly correlated (α = 0.05) pairwise positions in the orthogroup and eUSS alignments (total number of significant Φ*c* values divided by the length of the sequences in the orthogroup).

**Figure 8.**
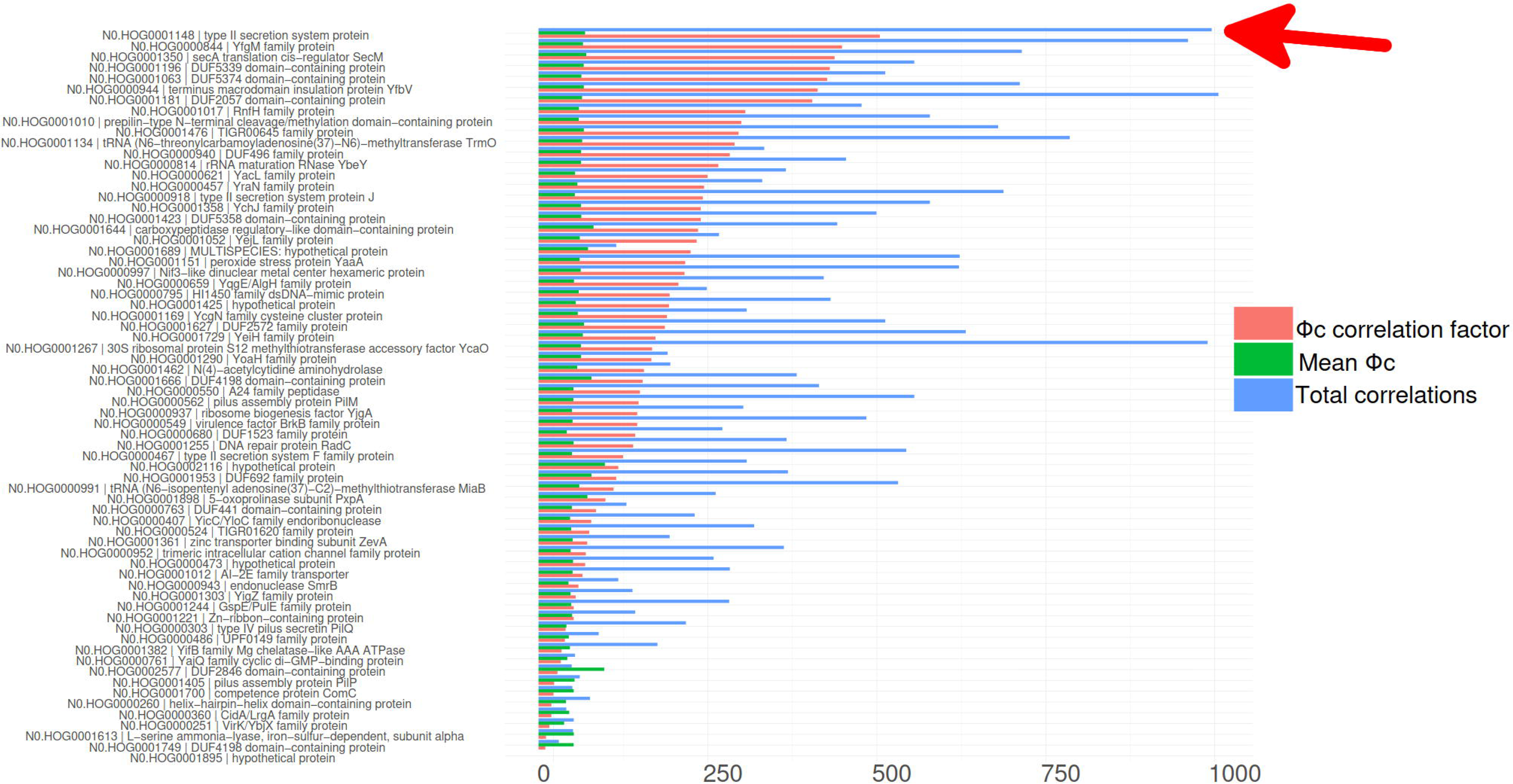
Coevolution of all modeled orthogroups and eUSS by Cramér’s Φ. Combined bar plots showing Φ*c* factor (red), mean Φ*c* (green) and total number of significant Φ*c* correlations (blue) for all orthogroups, sorted by mean Φ*c*. Φ*c* correlation factor and mean Φ*c* were multiplied by 100 to better scale with the number of Φ*c* correlations. Vertical axis labels: OrthoFinder hierarchical orthogroups and its most common gene name. Horizontal axis labels: Natural logarithm (ln). The relative positioning of PpdA = N0.HOG0001148 type II secretion system protein is indicated by red arrow.

PpdA also ranked second in terms of total number of significant Φ*c* correlations (996), only exceeded by DUF2057 domain-containing protein (1006) (Figure 8, Table S4). YfgM family protein (YfgM) also had many significant pairwise Φ*c* correlations (961) with eUSS and a high (4.49) Φ*c* correlation factor (Figure 8, Table S4). Other proteins with a combination of high correlation factor and number of significant correlations were SecA translation cis-regulator SecM (SecM) and terminus macrodomain insulation protein YfbV (Figure 8, Table S4).

The PpdA correlations to each eUSS position showed that the first eUSS position 1(A/T) was correlated with many positions throughout PpdA, having the strongest correlations with PpdA positions 91, 149 and 68 (Φ*c* = 0.939; Φ*c*= 0.927; Φ*c*=0.894) (Table S5; Fig. 9). eUSS Position 2(A) did not vary across sequences and thus had no significant pairwise correlations with PpdA positions. All three eUSS positions 3(A/C), 4(G/A) and 5 (T/A) were correlated with the same positions in PpdA and of equal magnitude. These three positions distinguish the 9-mer Hin-USS [w/ 3(A),4(G) and 5(T)) from Apl-USS (w/ 3(C), 4(A) and 5(A)]. Since these three positions also separate the Hin-USS and Apl-USS phylogenetic clades (21), the heatmap highlights that many of the amino acids across PpdA are co-evolved (drift and selection) with each specificity. Several strong PpdA correlations (Φ*c* = 1) were found with these three eUSS positions, for example positions 12, 116 and 174 in PpdA (Table S5). The 9-mer inner-core GCGG (eUSS positions 6(G), 7(C), 8(G) and 9(G)) and eUSS position 10(T) were invariable, as shown by the grey cells in Fig. 8. eUSS position 11(C/T/G/A) contained all four nucleotides, with C the most prevalently predicted, and was correlated with several PpdA positions (Table S5). The strongest correlations were with respective PpdA positions 187 (Φ*c*=0.850), 133 (Φ*c*=0.834) and 191 (Φ*c*=0.812). eUSS position 12(A/T/G) was predicted with predominantly A, less T and G, and had moderately high correlations with PpdA positions 48 (Φ*c*=0.702); 50 (Φ*c*=0.603) and 98 (Φ*c*=0.598) and low to moderate correlations with other positions (Table S5). eUSS position 13 (A/G/T) was predicted with A, less G and T, and had several correlations with PpdA positions, of which 50 (Φ*c*=0.801), 71 (Φ*c*=0.786) and 147 (Φ*c*=0.738) were the strongest. eUSS position 14(A/T) was predicted with near equal amounts of A and T and was strongly associated with the three PpdA positions 153 (Φ*c*=0.796), neighboring 152 (Φ*c*=0.789) and 125 (Φ*c*=0.773). eUSS positions 15(T) and 16(T) were invariable as shown by the grey cells in Fig. 8. eUSS position 17(T/C) was predicted with mostly T and a small number of C and were strongly correlated (Φ*c*=1) with the three PpdA positions 42, 147 and 170 (Table S5).

**Figure 9.**
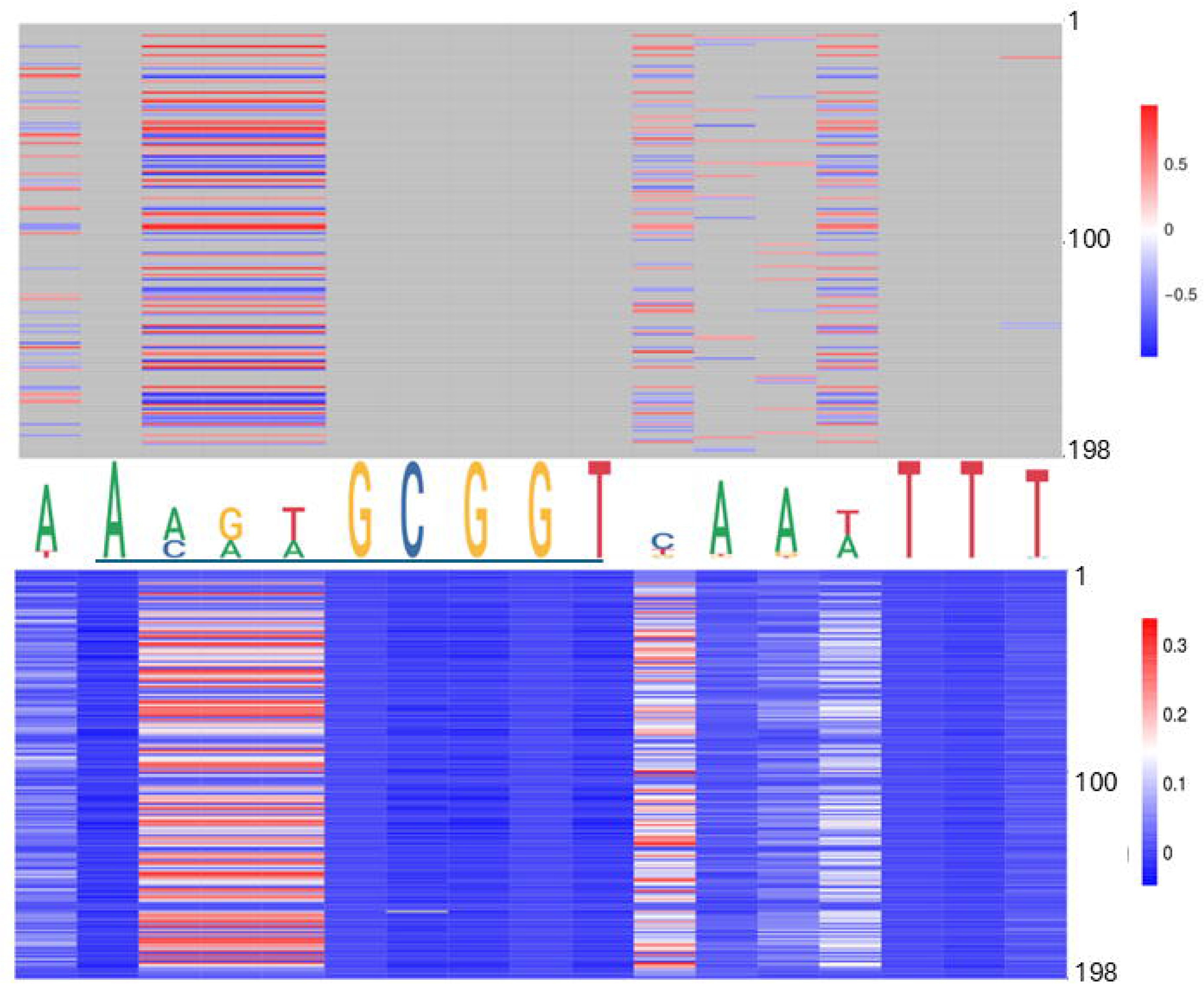
Significant pairwise correlations for PpdA positions and eUSS. PpdA positions on the vertical axis (1-197) and eUSS positions (1-17) on the horizontal axis as sequence logo with the 9-mer USS underlined. Upper panel: Φ*c* correlations according to given scale (-1 to +1), with pairwise positions without significant correlations in grey. Lower panel: miBIO correlations according to given scale (-0.5 to 0.37), with negative and non-correlating MI values in blue, neutral in white and positive in red.

Furthermore, PpdA was the second ranking orthogroup in the miBio coevolution analysis in regard of mean MI for positive MI values, surpassed only by tRNA N6-threonylcarbamoyladenosine(37)-N6)-methyltransferase TrmO (Table 1; Table S6; Fig. 10), ranking ninth in terms of MI factor (number of positive MI values divided by length of the sequences in the orthogroup). We noted that the *trmO* gene (Hi_0510) has two intragenic Hin-USS in *H. inf* Rd whereas *ppdA* (Hi_0938) has none (data not shown). Interestingly, PpdB, a protein predicted by STRING to be a functional partner of PpdA and also a pilin, ranked third in this analysis. YfgM family protein, which ranked second in the Φ*c* analysis, ranked fifth in MI factor. The pairwise sites that came out with the strongest positive MI values for PpdA-eUSS in miBIO showed considerable overlap with those in Φ*c* (Fig. 8-9, Tables S5 and S7), and the overall distribution of positive MI values and significant Φ*c* were highly similar. For eUSS position 1, two of the top three PpdA positions in the Φ*c* analysis were also among the top three correlating positions in the miBIO analysis, with the highest MI values being PpdA positions 25, 91 and 149, respectively (Tables S5 and S7). The invariable eUSS position 2(A) correlated with exactly three PpdA positions (16, 198 and 12) with positive MI values, albeit weakly positive. The USS dialect specific eUSS positions 3-5 correlated with PpdA position 12 with highest MI being the same position that ranked on top in Φ*c*, while multiple sites ranked second in both miBIO and Φ*c* such as PpdA position 116.

**Figure 10.**
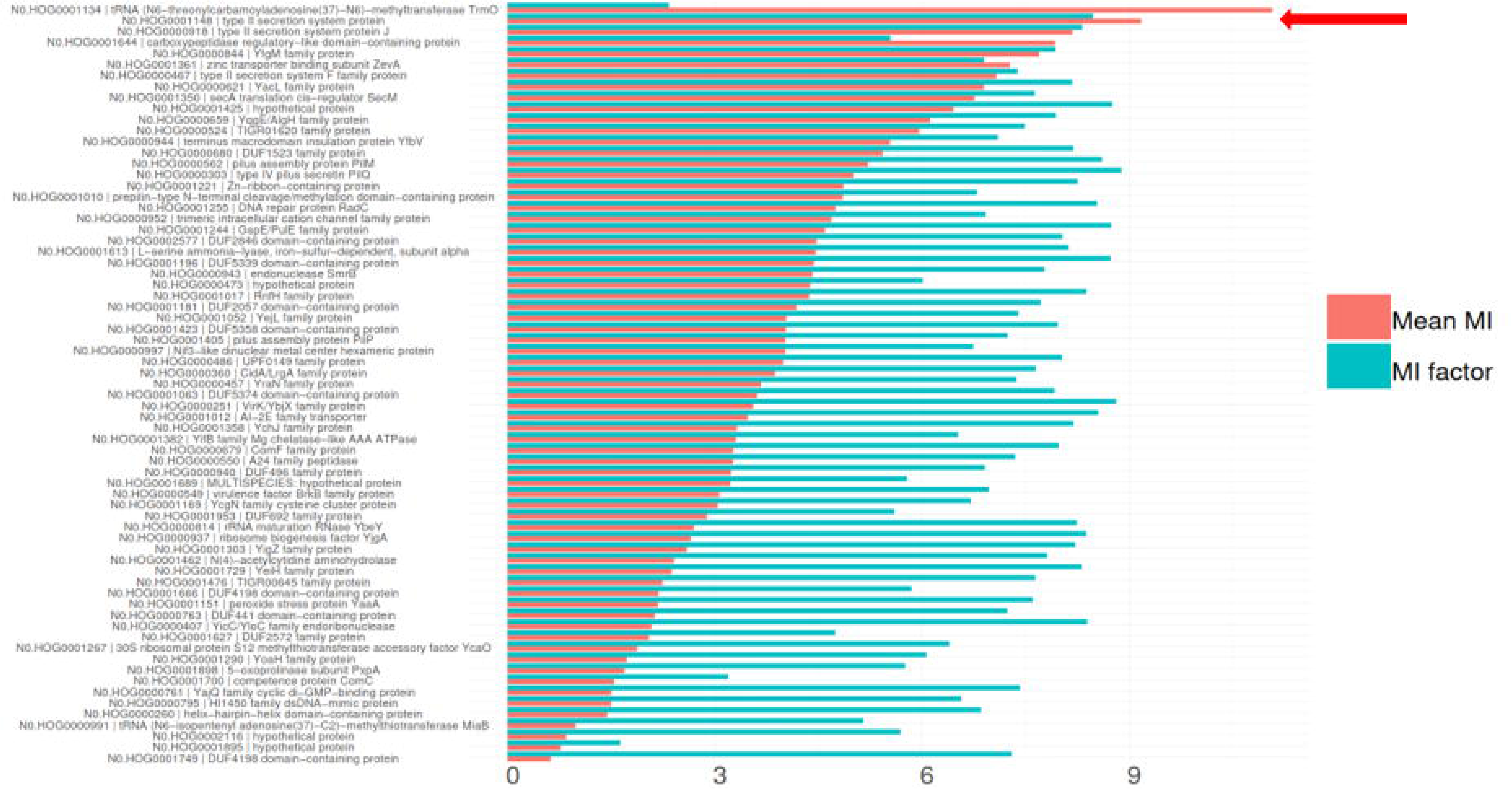
Coevolution of all modeled orthogroups and eUSS by miBIO. Combined bar plots showing mean MI and MI factor, sorted by mean MI. Mean MI was multiplied by 100 to better scale with MI factor. OrthoFinder hierarchical orthogroup numbers and gene names are listed. The relative positioning of PpdA = N0.HOG0001148 type II secretion system protein is indicated by red arrow.

**Table 1.** The top ten orthogroups in terms of mean MI in miBIO. Coverage refers to the number of OD species with orthologs in the orthogroup.

| OrthoFinder HOG | OrthoFinder OG | Gene name | OrthoFinder coverage | Protein length | Mean non-negative MI |
| --- | --- | --- | --- | --- | --- |
| N0.HOG0001<br>134 | OG0000950 | tRNA (N6-threonylcarbamoyladenosine(37)-N6)-methyltransferase TrmO | 147 | 272 | 0.1108185209<br>3349 |
| N0.HOG0001<br>148 | OG0000964 | type II secretion system protein | 147 | 198 | 0.0918067492<br>90887 |
| N0.HOG0000<br>918 | OG0000734 | type II secretion system protein J | 148 | 283 | 0.0818427395<br>35936 |
| N0.HOG0000<br>844 | OG0000660 | YfgM family protein | 149 | 215 | 0.0770323590<br>53076 |
| N0.HOG0001<br>361 | OG0001177 | zinc transporter binding subunit ZevA | 137 | 270 | 0.0727758381<br>93616 |
| N0.HOG0000<br>467 | OG0000283 | type II secretion system F family protein | 149 | 434 | 0.0708391122<br>89246 |
| N0.HOG0000<br>621 | OG0000437 | YacL family protein | 150 | 147 | 0.0690436856<br>56941 |
| N0.HOG0001<br>350 | OG0001166 | secA translation cis-regulator SecM | 141 | 164 | 0.0676582309<br>55378 |
| N0.HOG0000<br>659 | OG0000475 | YqgE/AlgH family protein | 149 | 204 | 0.0612235444<br>67119 |
| N0.HOG0000<br>524 | OG0000340 | TIGR01620 family protein | 146 | 422 | 0.0596713961<br>1252 |

In contrast to the Φ*c* analysis, miBIO results showed eUSS positions 6-10 to weakly correlate with different PpdA positions of which 7 and 16 stood out. eUSS position 11 had PpdA positions 191, 133 and 98 as the three with the highest MI, of which the two former were also among the top ranking three in Φ*c*. For eUSS position 12, PpdA positions 89, 47 and 117 ranked among the top three, respectively, while for eUSS position 13, PpdA positions 177, 175 and 71 ranked top three. eUSS position 14 had PpdA positions 151, 97 and 122 ranked the top three, while eUSS position 15 correlated with PpdA positions 7, 16 and 11 as the top three. For eUSS position 16, PpdA positions 7, 16 and 198 ranked the top three. eUSS position 17 correlated with several PpdA positions of which 42, 147 and 170 ranked first and among top three in the miBIO and Φ*c* analysis, respectively.

## Discussion

In searching for the USS-receptor, Pasteurellaceae orthologs were identified (eggNOG) using the *H. inf* Rd genome as reference. *H. inf* Rd was the first bacterial genome to be fully sequenced (47) and was the most fitting reference since almost all experimental uptake and transformation data are from this species and strain. The wild type phenotype of *H. inf* Rd displays very high transformation frequencies (11) documenting the presence of both an expressed and highly specific USS-binding protein. The 73O dataset was considered comprehensive in being based on rich GO annotations and functional terms known to be relevant for DNA uptake, transformation and competence. We expected that the Pasteurellaceae would have USS-specificity encoded by a surface exposed protein constituting a part of current transformation models (8) including that of *L. pneumophila* (25) which is also a γ-proteobacterium. It has also been firmly established in DNA binding and uptake studies that USS-binding takes place on the extracellular side of the outer membrane (12) warranting the focus on proteins and protein domains facing this environment. The proteins filtered out at the eggNOG/STRING stage were consequently excluded based on functional annotations and low orthogroup coverage, although in some cases, especially for DUF proteins and hypothetical proteins, the evidence for function and molecular interactions were scarce and only predicted. We consider all relevant proteins to be modeled by AF3 here. Although it is theoretically possible to model all OD orthogroups, this would be computationally exhaustive and would include many orthogroups that can not serve as the USS receptor due to functional annotations or low orthogroup coverage.

PpdA was the only orthogroup that consistently yielded AF3 models of high quality (high ipTM; low PAE; high pLDDT; low CPPM). Furthermore, PpdA was the only protein that yielded high quality models in all ten modeled species. Also, this orthogroup was the only for which orthogroup_USS_ models had significantly higher ipTM than orthogroup_scr_. Thus, we find that the AF3 results alone convincingly target PpdA as the USS receptor in Pasteurellaceae.

PpdA (HI0938) has previously been found to constitute part of the competence regulon in *H. inf* (48) under the name ComN. The first PpdA (HI0938/ComN) knock out was made in *H. inf* by Molnar (35) who observed loss of competence in this knock-out strain. This study could however not exclude that the competence loss in Δ*comN* was due to a polar effect on the downstream genes in the same operon (*comNOPQ*). This study also excluded ComE1 (HI1008) as the USS-receptor. Molnar (35) used an earlier version of the localization prediction algorithm DeepTMHMM (TMHMM v. 2.0) (49,50) to focus the USS-receptor search for cell surface exposed proteins (35). The limitations of the Δ*comN* strain was later amended using in-frame deletions of all genes known to be involved in competence, including *ppdA/comN*, and confirmed the loss of DNA uptake and transformation in the single *comN* mutant (18). PpdA (ComN) as part of the *comNOPQ* operon has for many years been one of several USS-receptor candidates based on gene regulatory and null-mutant phenotypes. Previous data (18,35) therefore align with our study arriving at PpdA as the predicted USS-receptor from candidate reducing efforts (function, interacting networks, localization and ortholog family coverage), deep-learning protein-DNA complex modeling, sequence specificity predictions and coevolutionary approaches. Molnar (35) characterized six other genes/proteins as plausible USS-specific candidates in addition to ComN included in our 73O dataset [HI0436 (*comD*); HI0438 (*comE*); HI0299 (*pilA*), HI0939 (*comO*) HI0940 (hypothetical), HI0941 (hypothetical)], yet none of our AF3 models of these were found to be within ipTM_r_. Although other proteins apart from PpdA yielded some high confidence models in AF3, these were either scarce, as illustrated by the distribution of ipTM values (see Fig. 3 A), or the scrambled models (orthogroup_scr_) for these proteins were of significantly higher quality than the native (orthogroup_USS_) ones, which was the case for N0.HOG0002116 and DUF4198-domain containing protein. The former, N0.HOG0002116, was found present only in 2/10 modeled and 44/150 OD species, thus it cannot likely serve as the USS receptor for all Pasteurellaceae species enriched in USS. The latter, DUF4198-domain containing protein_USS/scr,_ were modelled only for *H. inf* Rd and *A. actinomycetemcomitans* strain 31S (three DUF4198-domain containing protein_scr_ models), and thus poorly supported by AF3 as the USS receptor. Overall, the scarcity of good AF3 models for non-PpdA proteins, in contrast to the robust support for high-quality models for PpdA, supports PpdA as the USS receptor. Furthermore, PpdA shares topological and structural features with the two other well-characterized extracellular DNA-binding minor pilins ComP (24) and FimT (25) in Gram negative bacteria. PpdA resembles the DUS-specific ComP in having a disulfide-bridge anchored loop traversing the beta sheet. Since this anchored DD-loop in ComP is required for specific DUS-binding, it is tempting to speculate that the similar loop in PpdA fills a similar function in regard to USS-specificity. Further laboratory experiments are required to test this hypothesis. We and others (20,21,51) have previously suspected DUS and USS specificity to be an example of convergent evolution since the DUS/USS motifs are completely different, but identifying another minor pilin as the strongest USS-receptor candidate in Pasteurellaceae could suggest divergent evolution with deep roots in the Proteobacteria.

The results from AF3 and DeepPBS were in agreement in that AF3 consistently modeled PpdA_USS_ with high confidence while DeepPBS predicted specifically the two USSs as binding motifs. It is important to note that potential artifacts in the AF3 results are likely to have been echoed downstream by DeepPBS, as both are neural networks and have similarities in architecture (personal communication with DeepPBS developers). Yet, the convolutions in DeepPBS are applied to separate DNA (Sym-helix) and protein entities which potentially makes the DeepPBS results more independent from the robust AF3 models used. Although some overlap in models and predictions cannot fully be excluded, we consider these convergent results to be strong lines of support for PpdA being the USS receptor. Although AF3 robustly and uniquely modeled an alternative *F. canicola* strain HPA 21 PpdA_Apl-USS_ binding mode relative to the other AF3 PpdA_USS_ models (see Fig S2 J relative to Fig S2 A-I), DeepPBS predicted the similar binding specificity to closely match USS similar to the other predictions. This observation suggests that DeepPBS and AF3 build their respective protein-DNA complex models independently using separated DNA and protein input structures.

It is noteworthy that DeepPBS efficiently distinguished the USS dialect-specific nucleotides (AGT/CAA) yet predicts the GCGG-core common to both Hin- and Apl-USS less convincingly, as exemplified by the high mean probabilities for AGT and CAA in positions 2-4 for PpdA_Hin-USS_ and PpdA_Apl-USS_, respectively, as well as the strong prediction of **TT**GG in place of the consensus inner **GC**GG-core for PpdA_Apl-USS_ in terms of mean probability. It could be that there are elements of the DNA-binding mode of PpdA_USS_ AF3 models that accounts for this discrepancy/peculiarity. We speculate that the GCGG-core could be necessary for downstream catalytic activity or opening of the DNA-helix, while the AGT/CAA Hin/Apl differences accounts for the strongest specific PpdA-DNA binding and potentially the initial interaction. Molecular dynamics simulations could shed further light on these hypothesized different stages of the PpdA-USS interaction towards a fully resolved and stable complex. We include two videos (Supporting Videos 1 and 2) showing how the DeepPBS predictions change with the dynamically flexing PpdA-USS complex where individual frames will be analyzed in future work for co-varying nt predictions with dynamic amplitudes.

When comparing the DeepPBS predictions with the Hin-USS conservation as shown in the sequence logos we find informative cases where the weakest predicted positions were also the least conserved positions of the USS. The conservation of each USS position in the genome differs between species, suggesting differences in the specificity of their USS receptors. Examples where USS conservation and DeepPBS predictions correlate are the uniquely underrepresented 9(T) in *H. inf* Rd PpdA_USS_ predictions that matches the uniquely less conserved 9(T) in the *H. inf* sequence conservation logo. All other species’ PpdAs had considerably stronger 9(T) predictions that were also reflected in their USS conservation. The 9(T) of *P. multocida* USS, for example, is better conserved than both the two neighboring positions 7(G) and 8(G) and was predicted 9(T) with strong significance (Table S6). Another notable position is 3(G) that was the least significantly overrepresented predicted position in *M. succiniciproducens,* the only species predicted with significant overall 100% Hin-USS match, with the least conserved 3(G) of the Hin-USSs. Finding the sequence predictions PpdA_Hin-USS_ to match conservation at the species level further supports PpdA as the USS-receptor in these species.

In the Apl-USS group however, no apparent matches between conservation and predictions were observed. For example, 2(C) is better conserved in *F. canicola,* P. bacterium and *M.bovis* than in *A. lignieresii* and *A. equuli*, albeit this was also predicted significantly overrepresented in *A. equuli and A. lignieresii*. We speculate that the reason for not being able to identify similar patterns as in the Hin-USS group, is that the Apl-USS is considerably less conserved overall, with weaker conserved 1(A) and 2(C) and that the magnitude of their genomic enrichment is significantly lower (400>600/mb) than in the Hin-USS group (>800 Hin-USS/mb). Also, specific uptake of Apl-USS in *Actinobacillus pleuropneumoniae* (an Apl-USS species) has previously been shown to increase only 17-fold relative to scrambled USS which is in stark contrast to the >1500 fold increase in Hin-USS specific uptake in *H. influenzae* (21). These observations could suggest that USS specificity is considerably less pronounced in the Apl-USS group of species than in the Hin-USS group and explain our difficulties with identifying matches between the PpdA_Apl-USS_ DeepPBS predictions and Apl-USS conservation. Further laboratory experiments on Apl-USS specificity are needed to explore Apl-USS specificity in more detail and in a wider group of Pasteurellaceae bacteria with Apl-USS overrepresented.

Having found strong support for PpdA as the USS receptor from modeling and sequence specificity predictions, it does not seem coincidental that PpdA also had the highest correlation factor in the Cramér coevolution analyses of proteins and eUSS. The coevolutionary signal was higher by a Φ*c* correlation factor of 0.56 compared to YfgM family protein (SecYEG translocon-subunit), while PpdA came second to a methyltransferase (translation fidelity) in the miBIO analysis. A caveat with the coevolutionary analyses is that coevolutionary signals may be caused by genetic drift aligning with the phylogenetic relationships in the input sequences. While the shuffling method in miBIO corrects for much of the phylogenetic drift, it remains a challenge to assess impact on coevolution particularly since the two USS dialects adhere to phylogeny. It is also possible that proteins involved in the uptake process are imbued with coevolutionary relationships with (e)USS. We find for example several of the type IV pilus machinery components (PilM, PilP, PilQ, ComF and other minor pilins than PpdA) among the coevolved orthogroups showing these relationships. Also, due to the dependency between the USS-receptor and genomic enrichment of USS, several proteins would be in co-evolutionary relationships with the USS-receptor and hence also indirectly to (e)USS. The secA translation cis-regulator SecM and YfgM (SecYEG translocon subunit) with high correlation values in our analyses, are such examples in that all pilins involved in USS-uptake specifically and non-specifically are dependent on Sec machinery translocation (52). SecM has been shown to upregulate the functionality of SecA and the SecA pathway (51) to which *ppdA* and other pilins have adapted its prepilin signal peptide. Both the Φ*c* and miBIO results show co-evolution between eUSS and several of the other type II secretion system proteins which are necessary for USS-mediated transformation. Since PpdA has the highest coevolutionary scores of these Type IV pilus machinery components, it supports that PpdA as the USS-receptor relative to the other type II secretion system proteins generally and pilins specifically. An additional caveat is that regions of the genome with high eUSS content may disproportionately contribute to transformation, potentially generating spurious coevolutionary signals due to physical linkage between the eUSS and neighboring coding sequences or CDS rich in eUSS. The top-scoring miBIO TrmO may be an example of this co-evolutionary process since the *trmO* CDS is enriched with two eUSSs in contrast to none in *ppdA* itself. These signals may reflect transformation-driven linkage disequilibrium rather than true evolutionary coupling between residues or sites. In these cases, the partitioning of (e)Hin-USS and (e)Apl-USS will thus closely follow the phylogeny and cause an inflation of the coevolutionary signal between these proteins and eUSS. Further studies of USS enrichment could shed further on this mode of evolution (manuscript in prep.). All of these problems could result in falsely positively high Φ*c* and MI values in the coevolutionary analyses. However, we acknowledge that all proteins tested are faced with these challenges and thus the coevolution results are still helpful in pinpointing likely candidate USS receptor proteins. Perhaps even more so, the coevolution results tell us which proteins are likely *not* the USS receptor, as high coevolutionary signal would be expected for the true USS receptor having shaped USS-rich Pasteurellaceae genomes in deep time.

Combining the AF3 and DeepPBS results with the coevolution results for the AF3 and DeepPBS predicted USS receptor, shed some light on potential binding sites in the PpdA-(e)USS complex. We find that the eUSS-PpdA pairwise positions vary across the different modeled species leaving the elucidation of specific binding sites unresolved. A unified characterization of the binding interface from the AF3, DeepPBS and coevolution results would thus be a complex and integrative task for future research alongside wet-lab validation of PpdA as the USS-receptor. The PpdA-USS modeling data presented here showing interactions with both the major and minor grooves of USS could aid these efforts. We find a strong overall structural similarity between PpdA and the *L. pneumophila* DNA receptor FimT indicating that minor pilins with such folded globular domains function as DNA receptors. We hypothesize further that the particular β-sheet traversing C-terminal loop connected by disulfide bridges in both ComP and PpdA could represent a characteristic feature of minor pilins displaying DNA-binding specificity. Further studies could test this and potentially better inform the concept of convergent vs. divergent evolution of uptake specificity in these families and identify key events in the evolution of the deeply rooted transformation mechanism.

### Data availability

The code and data used in this study can be accessed at https://github.com/stianale/USS-receptor.

