## Supplementary material for "Type IV minor pilin ComN predicted the USS-receptor in Pasteurellaceae": Supporting Information Figures S1-S6.pdf

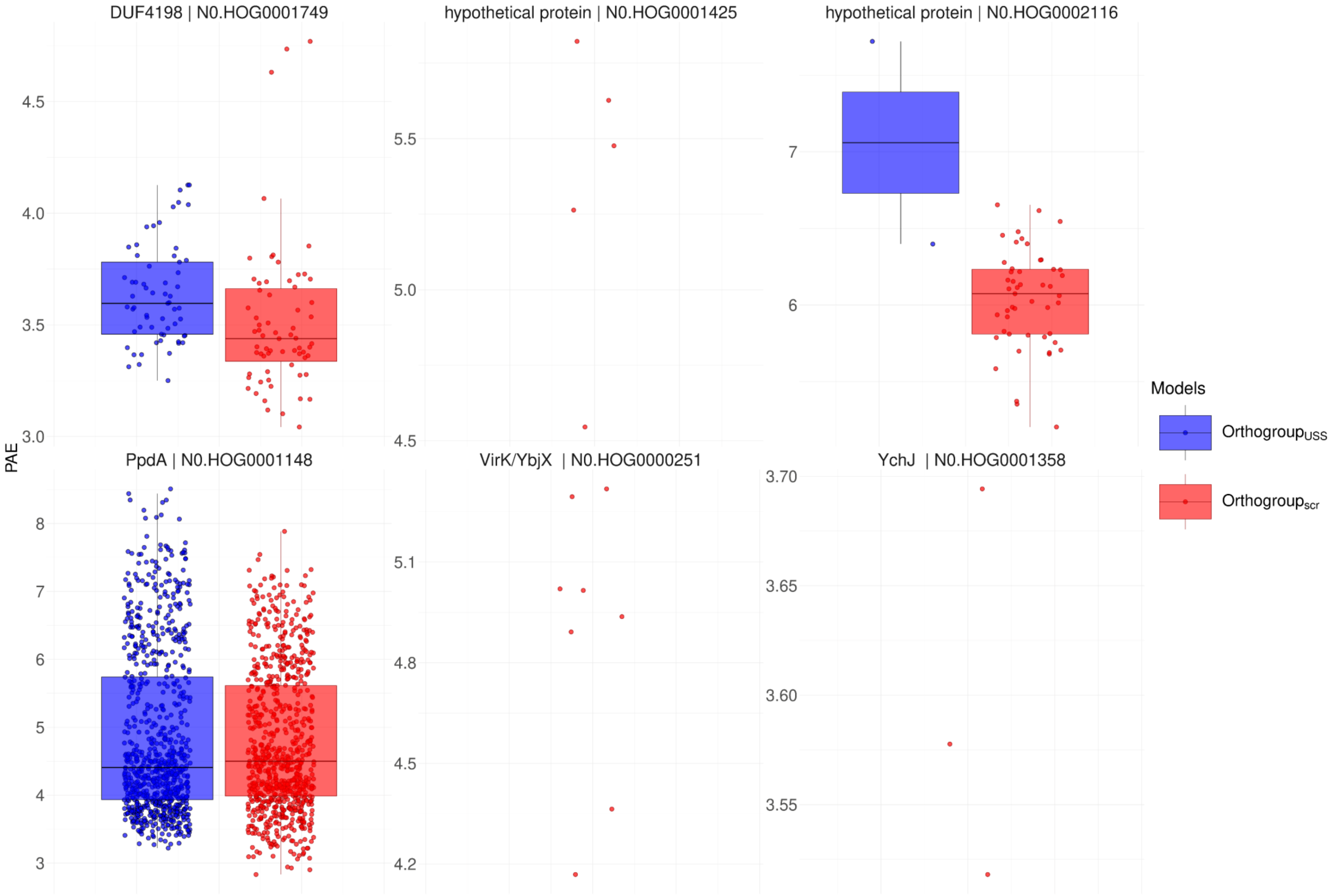


**Figure S1**. Box plots of distribution of PAE across all AF3 orthogroup$_{USS}$ and orthogroup$_{scr}$ models within ipTM$_{r}$ for the six modeled candidate proteins.

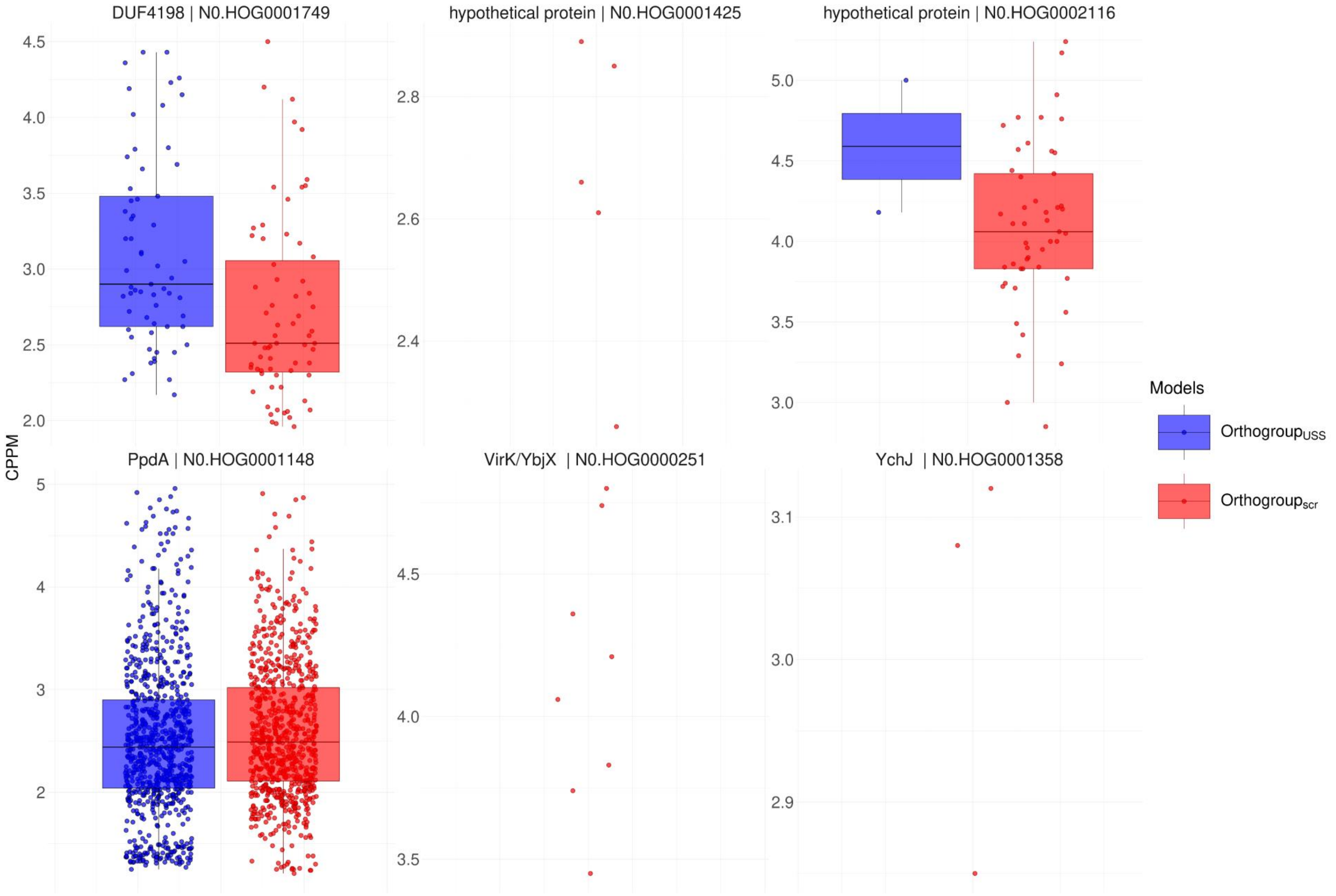


**Figure S2**. Box plots of distribution of CPPM across all AF3 orthogroup$_{USS}$ and orthogroup$_{scr}$ models within ipTM$_r$ for the six modeled candidate proteins.

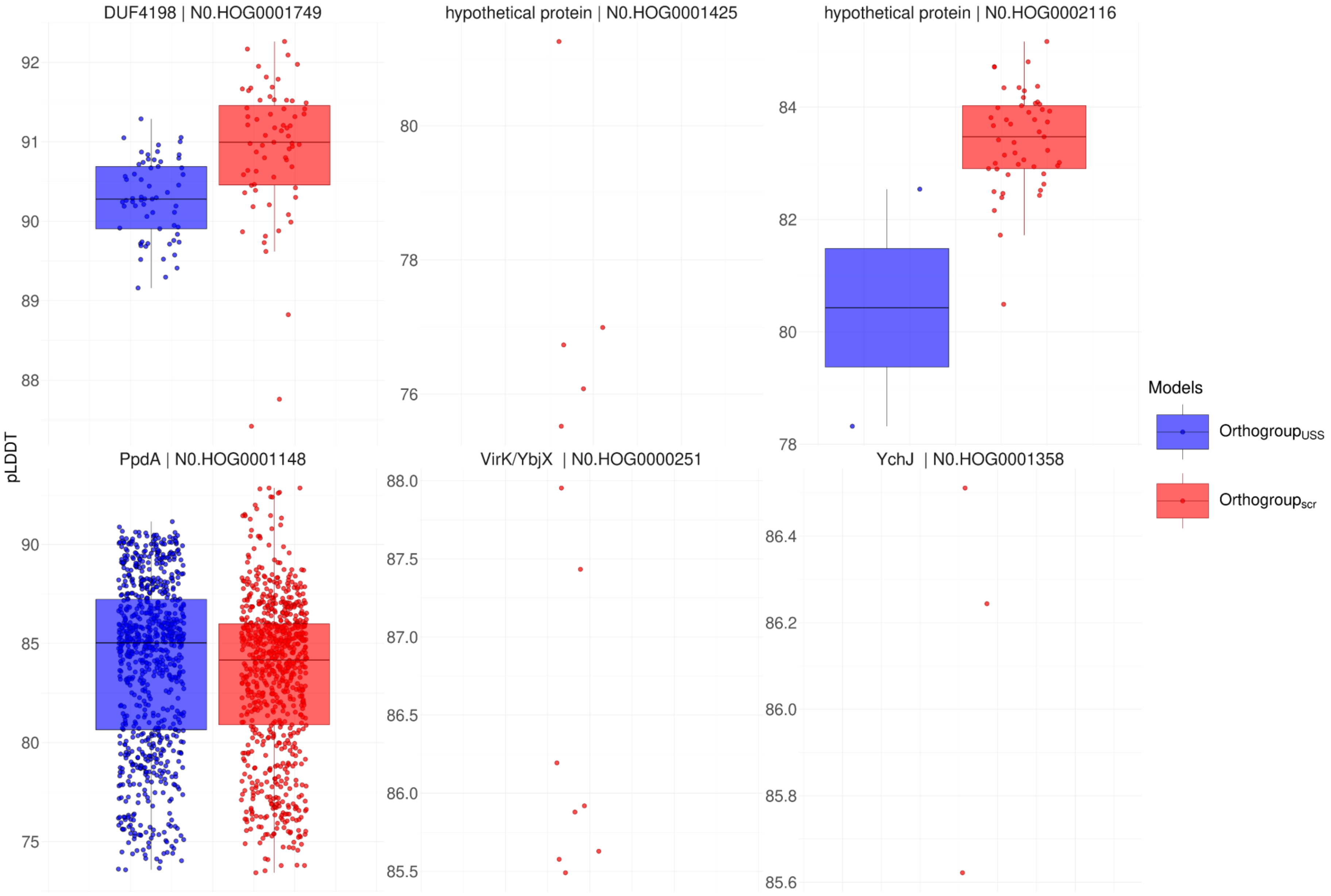


**Figure S3**. Box plots of distribution of pLDDT across all AF3 orthogroup$_{USS}$ and orthogroup$_{scr}$ models within ipTM$_r$ for the six modeled candidate proteins.

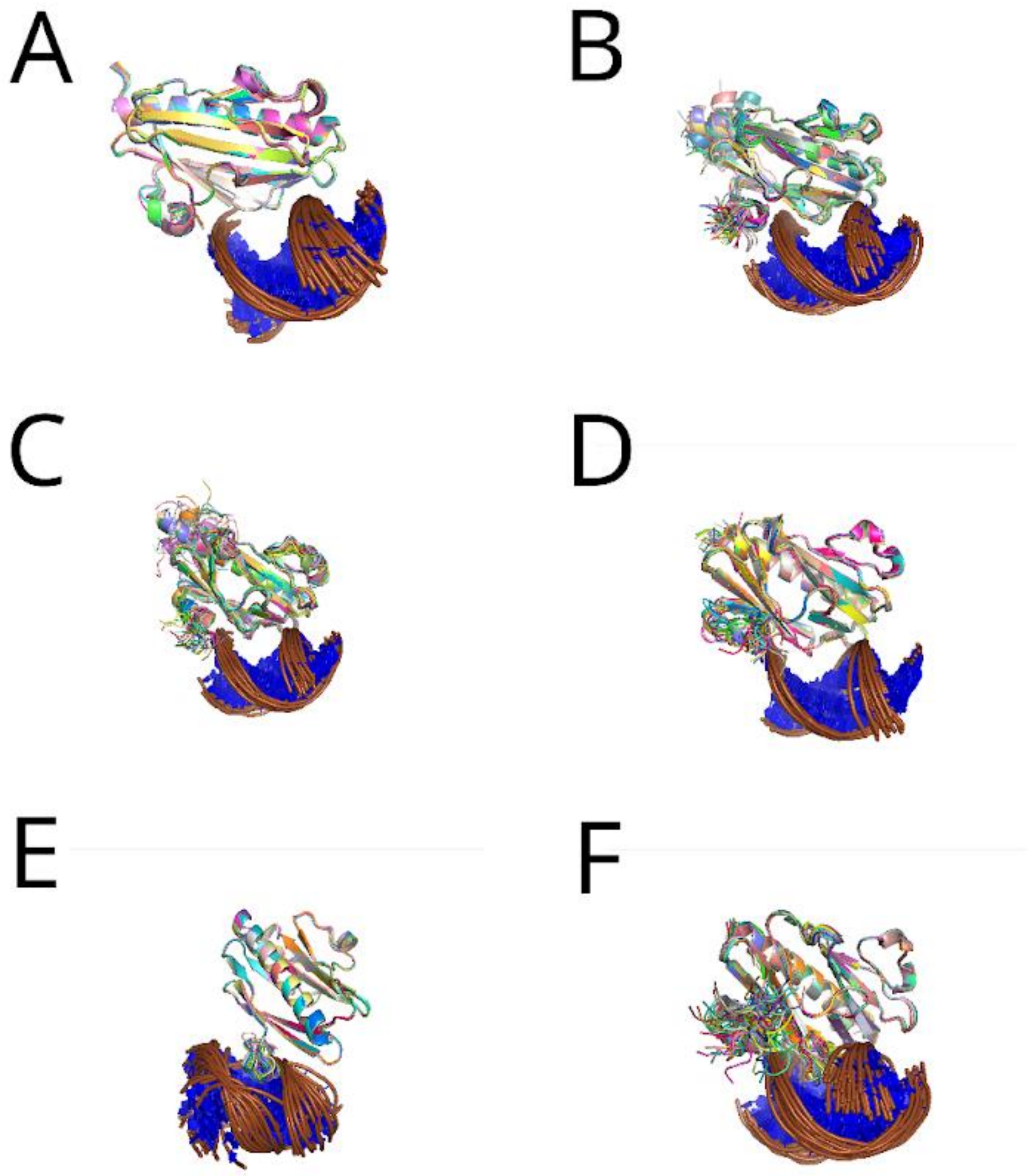


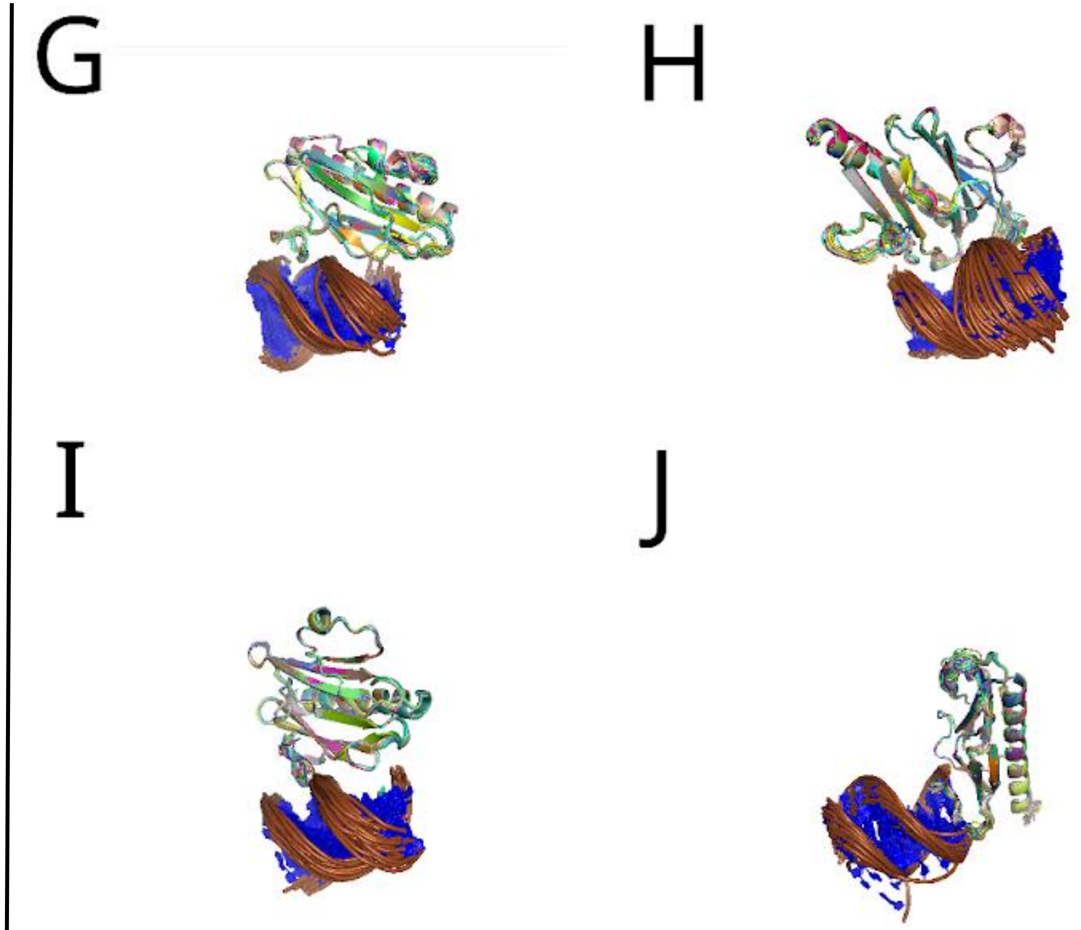


**Figure S4 Structural overlays of the largest clusters of PpdA$_{\mathrm{USS}}$ AF3 models with ipTM $\geq$ 0.6 and reciprocal DockQ $\geq$ 0.490.**
**A:** *Haemophilus influenzae* Rd. **B:** *Aggregatibacter sp* oral taxon 513.
**C:** *Aggregatibacter actinomycetemcomitans* strain 31S. **D:** *Pasteurella multocida* strain NCTC8282. **E:** *Pasteurellaceae bacterium* Orientalotternb1. **F:** *Mannheimia succiniciproducens* strain MBEL55E. **G:** *Mannheimia bovis* strain 39324S-11.
**H:** *Actinobacillus equuli subsp.* haemolyticus strain 3524. **I**: *Actinobacillus lignieresii* strain NCTC4189. **J**: *Frederiksenia canicola* strain HPA 2

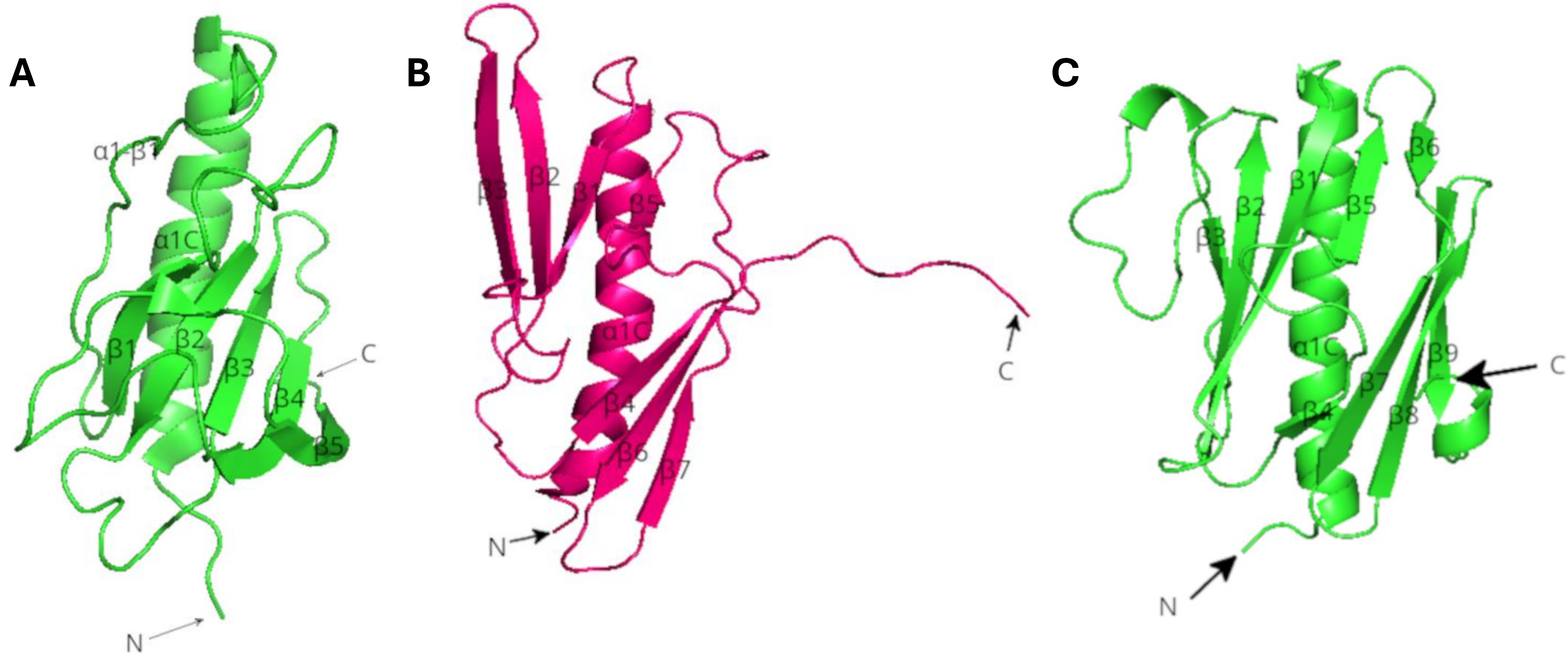


**Figure S5. 3D structures of validated minor pilin DNA receptors from different Proteobacteria and predicted PpdA.**
**A**: AF3-predicted structure of *Neisseria subflava* NJ9703$_{\mathrm{ComP}}$ from Helsem *et al.* (2025). **B**: NMR structure of *Legionella pneumophila* FimT (PDB: 7QYI). **C**: Top ranking *H. inf* Rd PpdA$_{\mathrm{Hin\text{-}USS}}$ model from AF3 (DNA masked).

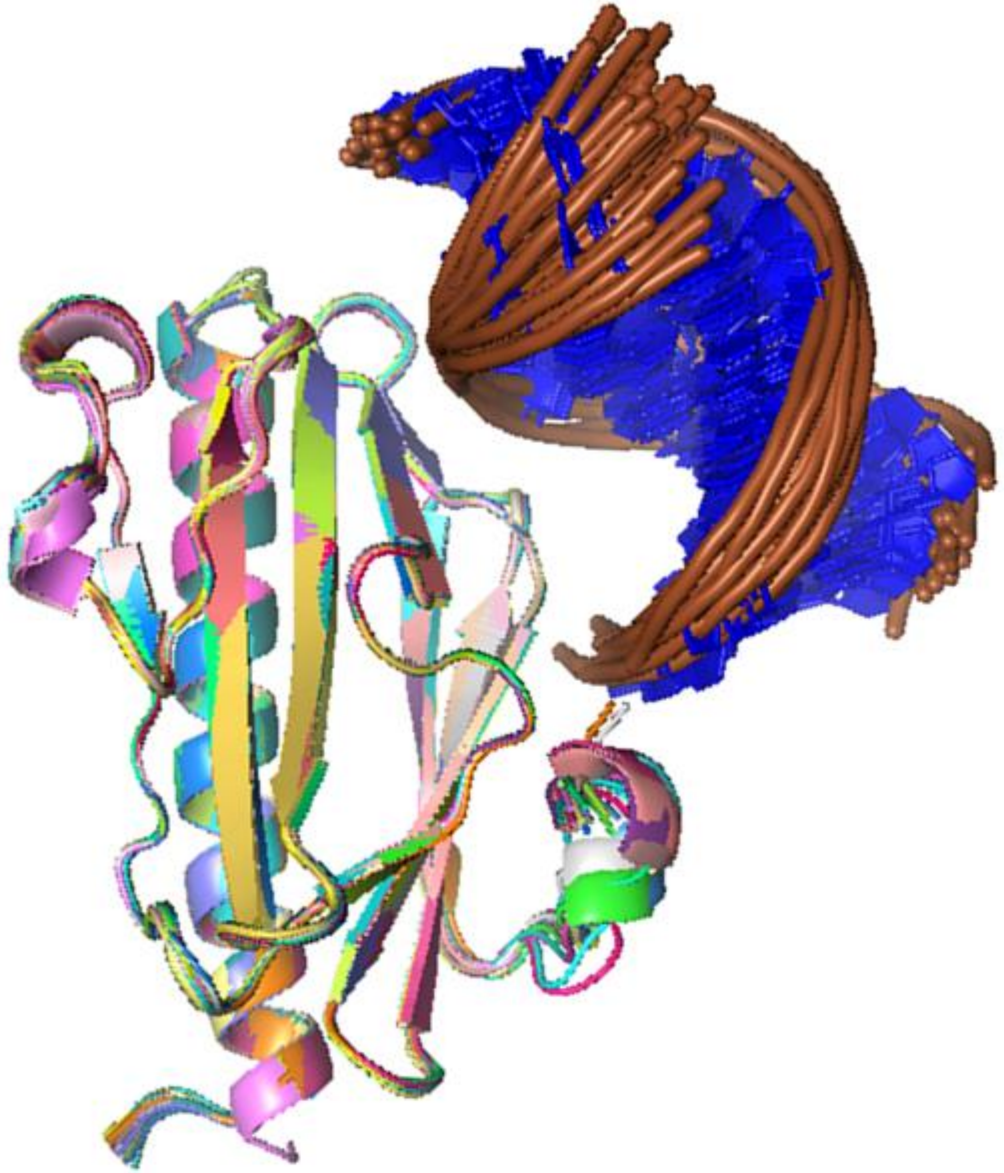

**Figure S6. The PpdA-USS AF3 predicted binding mode.**
The largest cluster of *H. inf* Rd $PpdA_{Hin\text{-}USS}$ models with reciprocal DockQ $\geq$ 0.49 aligned on the α-carbons of the top-ranking model.
