## Supplementary material for "Type IV minor pilin ComN predicted the USS-receptor in Pasteurellaceae": Supporting Information_legends and statistics.pdf

No. of supporting Figures: 6

No. of supporting Videos: 2

No. of supporting Tables: 9

No. of supporting data: 1

No. of supporting Code output: 1

### Supplementary information legends and statistical analysis

**Video S1.** DeepPBS predicted DNA-binding specificity for H. inf Rd 98/200 ppd$A_{Hin-USS}$ AF3 models with ipTM $\geq$ 0.6 (number of frames/models) concatenated into GIF format. Upper left panel: AF3 input Hin-USS sequence. Bottom left panel: DeepPBS prediction. Right panel: Sequence logos showing the relative proportions of the nucleotides at each position. Position 0 refers to USS position 1.

**Video S2.** DeepPBS predicted DNA-binding specificity for 98/200 *Actinobacillus equuli* ppd$A_{Apl-USS}$ AF3 models with ipTM $\geq$ 0.6 concatenated into GIF format. Upper left panel: AF3 input Apl-USS sequence. Bottom left panel: DeepPBS prediction. Right panel: Sequence logos showing the relative proportions of the nucleotides at each position.

**Supplementary Data 1** PpdA alignment across all 150 OD species generated using MView v. 1.68 (1)

**Supplementary Code Output S1** Wilcoxon rank sum tests on ipTM, PAE, CPPM and pLDDT (orthogroup$_{USS}$ vs. orthogroup$_{scr}$).

```
> results <- new_df %>%
+   group_by(Protein_HOG) %>%
+   filter(n_distinct(Scrambled) > 1) %>%  # Ensure both "Yes" and "No" are present
+   summarise(p_value = list(wilcox.test(ipTM ~ Scrambled)$p.value), .groups = "drop") %>%
+   unnest(p_value)
Warning message:
There was 1 warning in `summarise()`.
ℹ In argument: `p_value = list(wilcox.test(ipTM ~ Scrambled)$p.value)`.
ℹ In group 2: `Protein_HOG = "hypothetical protein | N0.HOG0002116"`.
Caused by warning in `wilcox.test.default()`:
! cannot compute exact p-value with ties
>
> print(results)
# A tibble: 3 × 2
  Protein_HOG                                         p_value
  <chr>                                                 <dbl>
1 DUF4198 domain-containing protein | N0.HOG0001749 0.0000156
2 hypothetical protein | N0.HOG0002116              0.178
3 type II secretion system protein | N0.HOG0001148  0.000000243
>
> results <- new_df %>%
+   group_by(Protein_HOG) %>%
+   filter(n_distinct(Scrambled) > 1) %>%  # Ensure both "Yes" and "No" are present
+   summarise(p_value = list(wilcox.test(PAE ~ Scrambled)$p.value), .groups = "drop") %>%
+   unnest(p_value)
>
> print(results)
# A tibble: 3 × 2
```

```
  Protein_HOG                                          p_value
  <chr>                                                <dbl>
1 DUF4198 domain-containing protein | N0.HOG0001749 0.000226
2 hypothetical protein | N0.HOG0002116              0.0392
3 type II secretion system protein | N0.HOG0001148  0.808
>
> results <- new_df %>%
+   group_by(Protein_HOG) %>%
+   filter(n_distinct(Scrambled) > 1) %>%  # Ensure both "Yes" and "No" are present
+   summarise(p_value = list(wilcox.test(CPPM ~ Scrambled)$p.value), .groups = "drop") %>%
+   unnest(p_value)
Warning message:
There was 1 warning in `summarise()`.
ℹ In argument: `p_value = list(wilcox.test(CPPM ~ Scrambled)$p.value)`.
ℹ In group 2: `Protein_HOG = "hypothetical protein | N0.HOG0002116"`.
Caused by warning in `wilcox.test.default()`:
! cannot compute exact p-value with ties
>
> print(results)
# A tibble: 3 × 2
  Protein_HOG                                          p_value
  <chr>                                                <dbl>
1 DUF4198 domain-containing protein | N0.HOG0001749 0.000119
2 hypothetical protein | N0.HOG0002116              0.190
3 type II secretion system protein | N0.HOG0001148  0.0107
>
> results <- new_df %>%
+   group_by(Protein_HOG) %>%
+   filter(n_distinct(Scrambled) > 1) %>%  # Ensure both "Yes" and "No" are present
+   summarise(p_value = list(wilcox.test(pLDDT ~ Scrambled)$p.value), .groups = "drop") %>%
+   unnest(p_value)
>
> print(results)
# A tibble: 3 × 2
  Protein_HOG                                            p_value
  <chr>                                                  <dbl>
1 DUF4198 domain-containing protein | N0.HOG0001749 0.0000000551
2 hypothetical protein | N0.HOG0002116              0.0392
3 type II secretion system protein | N0.HOG0001148  0.0000636
```
