## Supplementary material for "Type IV minor pilin ComN predicted the USS-receptor in Pasteurellaceae": Table S1.docx

**Table S1**	Gene Ontology (GO) terms and functional annotation inclusion criteria used to filter orthogroups in the eggNOG-mapper results.

| **Search term** | **Type** | **Description** |
| --- | --- | --- |
| GO:0009289 | GO evidence | Pilus |
| GO:0044096 | GO evidence | Type IV pilus |
| GO:0009297 | GO evidence | Pilus assembly |
| GO:0043683 | GO evidence | Type IV pilus assembly |
| GO:0140621 | GO evidence | Type 1 pilus |
| GO:0009986 | GO evidence | Cell surface |
| GO:0019867 | GO evidence | Outer membrane |
| GO:0005576 | GO evidence | Extracellular region |
| GO:0005620 | GO evidence | Periplasm |
| \tcompetence | Functional annotation | - |
| \tCompetence | Functional annotation | - |
| competence | Functional annotation | - |
| \tpil | Functional annotation | - |
| \tcom | Functional annotation | - |
| \tCom | Functional annotation | - |
| \tPil | Functional annotation | - |
| pilus | Functional annotation | - |
| pilin | Functional annotation | - |
| type II | Functional annotation | - |
| type IV | Functional annotation | - |
| Type II | Functional annotation | - |
| Type IV | Functional annotation | - |
| cleavage | Functional annotation | - |
| N-terminal | Functional annotation | - |
| peptidase | Functional annotation | - |
| hypothetical protein | Functional annotation | - |
| DUF | Functional annotation | - |
| UPF | Functional annotation | - |
