## Supplementary material for "Type IV minor pilin ComN predicted the USS-receptor in Pasteurellaceae": Table S4.docx

**Table S4	Condensed results for the Cramér’s V (Φ*c*) analysis for all orthogroups tested.** PpdA (grey highlight) is N0.HOG0001148 type II secretion system protein according to its OthoFinder classification.

| **OrthoFinder HOG** | **OrthoFinder OG** | **Gene name** | **OrthoFinder coverage** | **Protein lengths** | **Num. significant** *Φ_c_* | **Correlation factor** |
| --- | --- | --- | --- | --- | --- | --- |
| N0.HOG0001148 | OG0000964 | type II secretion system protein | 147 | 197 | 996 | 5.05 |
| N0.HOG0000844 | OG0000660 | YfgM family protein | 149 | 214 | 961 | 4.49 |
| N0.HOG0001350 | OG0001166 | secA translation cis-regulator SecM | 141 | 163 | 715 | 4.38 |
| N0.HOG0001196 | OG0001012 | DUF5339 domain-containing protein | 146 | 129 | 556 | 4.31 |
| N0.HOG0001063 | OG0000879 | DUF5374 domain-containing protein | 148 | 120 | 513 | 4.27 |
| N0.HOG0000944 | OG0000760 | terminus macrodomain insulation protein YfbV | 149 | 172 | 712 | 4.13 |
| N0.HOG0001181 | OG0000997 | DUF2057 domain-containing protein | 147 | 248 | 1006 | 4.05 |
| N0.HOG0001017 | OG0000833 | RnfH family protein | 148 | 156 | 478 | 3.06 |
| N0.HOG0001010 | OG0000826 | prepilin-type N-terminal cleavage/methylation domain-containing protein | 148 | 193 | 579 | 3 |
| N0.HOG0001476 | OG0001280 | TIGR00645 family protein | 121 | 229 | 680 | 2.96 |
| N0.HOG0001134 | OG0000950 | tRNA (N6-threonylcarbamoyladenosine(37)-N6)-methyltransferase TrmO | 147 | 271 | 786 | 2.9 |
| N0.HOG0000940 | OG0000756 | DUF496 family protein | 149 | 118 | 334 | 2.83 |
| N0.HOG0000814 | OG0000630 | rRNA maturation RNase YbeY | 149 | 171 | 455 | 2.66 |
| N0.HOG0000621 | OG0000437 | YacL family protein | 150 | 146 | 366 | 2.5 |
| N0.HOG0000457 | OG0000273 | YraN family protein | 147 | 135 | 331 | 2.45 |
| N0.HOG0000918 | OG0000734 | type II secretion system protein J | 148 | 282 | 688 | 2.43 |
| N0.HOG0001423 | OG0001236 | DUF5358 domain-containing protein | 131 | 208 | 500 | 2.4 |
| N0.HOG0001358 | OG0001174 | YchJ family protein | 140 | 241 | 579 | 2.4 |
| N0.HOG0001644 | OG0001442 | carboxypeptidase regulatory-like domain-containing protein | 93 | 187 | 442 | 2.36 |
| N0.HOG0001052 | OG0000868 | YejL family protein | 148 | 114 | 267 | 2.34 |
| N0.HOG0001689 | OG0001482 | MULTISPECIES: hypothetical protein | 88 | 51 | 115 | 2.25 |
| N0.HOG0001151 | OG0000967 | peroxide stress protein YaaA | 147 | 286 | 623 | 2.17 |
| N0.HOG0000997 | OG0000813 | Nif3-like dinuclear metal center hexameric protein | 148 | 287 | 622 | 2.16 |
| N0.HOG0000659 | OG0000475 | YqgE/AlgH family protein | 149 | 203 | 422 | 2.07 |
| N0.HOG0000795 | OG0000611 | HI1450 family dsDNA-mimic protein | 150 | 128 | 249 | 1.94 |
| N0.HOG0001425 | OG0001238 | hypothetical protein | 126 | 223 | 432 | 1.93 |
| N0.HOG0001169 | OG0000985 | YcgN family cysteine cluster protein | 147 | 162 | 308 | 1.9 |
| N0.HOG0001627 | OG0001425 | DUF2572 family protein | 96 | 273 | 513 | 1.87 |
| N0.HOG0001729 | OG0001522 | YeiH family protein | 82 | 364 | 632 | 1.73 |
| N0.HOG0001267 | OG0001083 | 30S ribosomal protein S12 methylthiotransferase accessory factor YcaO | 145 | 588 | 990 | 1.68 |
| N0.HOG0001290 | OG0001106 | YoaH family protein | 144 | 114 | 191 | 1.67 |
| N0.HOG0001462 | OG0001266 | N(4)-acetylcytidine aminohydrolase | 122 | 125 | 195 | 1.56 |
| N0.HOG0001666 | OG0001459 | DUF4198 domain-containing protein | 91 | 247 | 382 | 1.54 |
| N0.HOG0000550 | OG0000366 | A24 family peptidase | 148 | 276 | 415 | 1.5 |
| N0.HOG0000937 | OG0000753 | ribosome biogenesis factor YjgA | 148 | 207 | 303 | 1.46 |
| N0.HOG0000549 | OG0000365 | virulence factor BrkB family protein | 150 | 332 | 485 | 1.46 |
| N0.HOG0000680 | OG0000496 | DUF1523 family protein | 149 | 190 | 272 | 1.43 |
| N0.HOG0001255 | OG0001071 | DNA repair protein RadC | 144 | 261 | 367 | 1.4 |
| N0.HOG0000467 | OG0000283 | type II secretion system F family protein | 149 | 433 | 544 | 1.25 |
| N0.HOG0002116 | OG0001894 | hypothetical protein | 44 | 260 | 308 | 1.18 |
| N0.HOG0001953 | OG0001731 | DUF692 family protein | 56 | 320 | 369 | 1.15 |
| N0.HOG0000991 | OG0000807 | tRNA (N6-isopentenyl adenosine(37)-C2)-methylthiotransferase MiaB | 148 | 477 | 532 | 1.11 |
| N0.HOG0001898 | OG0001680 | 5-oxoprolinase subunit PxpA | 62 | 264 | 262 | 0.99 |
| N0.HOG0000763 | OG0000579 | DUF441 domain-containing protein | 148 | 152 | 130 | 0.85 |
| N0.HOG0000407 | OG0000226 | YicC/YloC family endoribonuclease | 150 | 293 | 231 | 0.78 |
| N0.HOG0000524 | OG0000340 | TIGR01620 family protein | 146 | 421 | 319 | 0.75 |
| N0.HOG0001361 | OG0001177 | zinc transporter binding subunit ZevA | 137 | 269 | 194 | 0.72 |
| N0.HOG0000952 | OG0000768 | trimeric intracellular cation channel family protein | 144 | 514 | 363 | 0.7 |
| N0.HOG0000473 | OG0000289 | hypothetical protein | 150 | 371 | 259 | 0.69 |
| N0.HOG0001012 | OG0000828 | AI-2E family transporter | 144 | 432 | 283 | 0.65 |
| N0.HOG0000943 | OG0000759 | endonuclease SmrB | 147 | 197 | 118 | 0.59 |
| N0.HOG0001303 | OG0001119 | YigZ family protein | 140 | 249 | 139 | 0.55 |
| N0.HOG0000303 | OG0000138 | type IV pilus secretin PilQ | 148 | 534 | 218 | 0.4 |
| N0.HOG0000486 | OG0000302 | UPF0149 family protein | 150 | 224 | 89 | 0.39 |
| N0.HOG0000761 | OG0000577 | YajQ family cyclic di-GMP-binding protein | 149 | 163 | 54 | 0.33 |
| N0.HOG0002577 | OG0002344 | DUF2846 domain-containing protein | 22 | 170 | 49 | 0.28 |
| N0.HOG0001405 | OG0001221 | pilus assembly protein PilP | 131 | 263 | 61 | 0.23 |
| N0.HOG0001700 | OG0001493 | competence protein ComC | 85 | 222 | 50 | 0.22 |
| N0.HOG0000360 | OG0000180 | CidA/LrgA family protein | 143 | 215 | 41 | 0.19 |
| N0.HOG0001613 | OG0001411 | L-serine ammonia-lyase, iron-sulfur-dependent, subunit alpha | 95 | 439 | 51 | 0.11 |
| N0.HOG0001749 | OG0001542 | DUF4198 domain-containing protein | 77 | 289 | 30 | 0.1 |
| N0.HOG0001895 | OG0001677 | hypothetical protein | 61 | 235 | 0 | 0 |
| N0.HOG0000251 | OG0000101 | VirK/YbjX family protein | 142 | 318 | 52 | 0.16 |
| N0.HOG0000562 | OG0000378 | pilus assembly protein PilM | 149 | 375 | 556 | 1.48 |
| N0.HOG0001221 | OG0001037 | Zn-ribbon-containing protein | 142 | 271 | 143 | 0.52 |
| N0.HOG0001244 | OG0001060 | GspE/PulE family protein | 143 | 534 | 282 | 0.52 |
| N0.HOG0001382 | OG0001198 | YifB family Mg chelatase-like AAA ATPase | 134 | 512 | 176 | 0.34 |
| N0.HOG0000260 | OG0000107 | helix-hairpin-helix domain-containing protein | 149 | 388 | 76 | 0.19 |
| N0.HOG0000679 | OG0000495 | ComF family protein | 148 | 260 | 146 | 0.56 |
