## Supplementary material for "Type IV minor pilin ComN predicted the USS-receptor in Pasteurellaceae": Table S6.docx

**Table S6**	**Condensed results from miBIO for all orthogroups tested.** OrthoFinder (hierarchical) orthogroups are listed, along with majority-vote gene name, coverage, protein length and mean MI for non-negative MIs. PpdA (grey highlight) is N0.HOG0001148 type II secretion system protein according to its OrthoFinder classification.

| **OrthoFinder HOG** | **OrthoFinder OG** | **Gene name** | **OrthoFinder coverage** | **Protein lengths** | **Mean non-negative MI** |
| --- | --- | --- | --- | --- | --- |
| N0.HOG0001134 | OG0000950 | tRNA (N6-threonylcarbamoyladenosine(37)-N6)-methyltransferase TrmO | 147 | 272 | 0.11081852093349 |
| N0.HOG0001148 | OG0000964 | type II secretion system protein | 147 | 198 | 0.091806749290887 |
| N0.HOG0000918 | OG0000734 | type II secretion system protein J | 148 | 283 | 0.081842739535936 |
| N0.HOG0000844 | OG0000660 | YfgM family protein | 149 | 215 | 0.077032359053076 |
| N0.HOG0001361 | OG0001177 | zinc transporter binding subunit ZevA | 137 | 270 | 0.072775838193616 |
| N0.HOG0000467 | OG0000283 | type II secretion system F family protein | 149 | 434 | 0.070839112289246 |
| N0.HOG0000621 | OG0000437 | YacL family protein | 150 | 147 | 0.069043685656941 |
| N0.HOG0001350 | OG0001166 | secA translation cis-regulator SecM | 141 | 164 | 0.067658230955378 |
| N0.HOG0000659 | OG0000475 | YqgE/AlgH family protein | 149 | 204 | 0.061223544467119 |
| N0.HOG0000524 | OG0000340 | TIGR01620 family protein | 146 | 422 | 0.05967139611252 |
| N0.HOG0000944 | OG0000760 | terminus macrodomain insulation protein YfbV | 149 | 173 | 0.055444329860066 |
| N0.HOG0000680 | OG0000496 | DUF1523 family protein | 149 | 191 | 0.054363729438059 |
| N0.HOG0000562 | OG0000378 | pilus assembly protein PilM | 149 | 376 | 0.052236226272417 |
| N0.HOG0000303 | OG0000138 | type IV pilus secretin PilQ | 148 | 534 | 0.050142796847737 |
| N0.HOG0001221 | OG0001037 | Zn-ribbon-containing protein | 142 | 272 | 0.048671258264436 |
| N0.HOG0001010 | OG0000826 | prepilin-type N-terminal cleavage/methylation domain-containing protein | 148 | 194 | 0.048571206702121 |
| N0.HOG0001255 | OG0001071 | DNA repair protein RadC | 144 | 262 | 0.047545773051832 |
| N0.HOG0000952 | OG0000768 | trimeric intracellular cation channel family protein | 144 | 514 | 0.046951264431508 |
| N0.HOG0001244 | OG0001060 | GspE/PulE family protein | 143 | 534 | 0.046013897025134 |
| N0.HOG0001196 | OG0001012 | DUF5339 domain-containing protein | 146 | 130 | 0.044380762425632 |
| N0.HOG0000943 | OG0000759 | endonuclease SmrB | 147 | 198 | 0.044208779182403 |
| N0.HOG0000473 | OG0000289 | hypothetical protein | 150 | 372 | 0.043863281196267 |
| N0.HOG0001017 | OG0000833 | RnfH family protein | 148 | 157 | 0.043717573768261 |
| N0.HOG0001181 | OG0000997 | DUF2057 domain-containing protein | 147 | 249 | 0.041881609714776 |
| N0.HOG0001052 | OG0000868 | YejL family protein | 148 | 115 | 0.04039686663819 |
| N0.HOG0000997 | OG0000813 | Nif3-like dinuclear metal center hexameric protein | 148 | 288 | 0.040266521078909 |
| N0.HOG0000486 | OG0000302 | UPF0149 family protein | 150 | 225 | 0.039962762897611 |
| N0.HOG0000360 | OG0000180 | CidA/LrgA family protein | 143 | 216 | 0.038751285501843 |
| N0.HOG0000457 | OG0000273 | YraN family protein | 147 | 136 | 0.036702936053794 |
| N0.HOG0001063 | OG0000879 | DUF5374 domain-containing protein | 148 | 121 | 0.036160750584191 |
| N0.HOG0000251 | OG0000101 | VirK/YbjX family protein | 142 | 319 | 0.035622829211873 |
| N0.HOG0001012 | OG0000828 | AI-2E family transporter | 144 | 433 | 0.034838243272496 |
| N0.HOG0001358 | OG0001174 | YchJ family protein | 140 | 242 | 0.033279433374233 |
| N0.HOG0001382 | OG0001198 | YifB family Mg chelatase-like AAA ATPase | 134 | 512 | 0.033095968088457 |
| N0.HOG0000550 | OG0000366 | A24 family peptidase | 148 | 277 | 0.032671986247792 |
| N0.HOG0000940 | OG0000756 | DUF496 family protein | 149 | 119 | 0.032371613522408 |
| N0.HOG0000549 | OG0000365 | virulence factor BrkB family protein | 150 | 333 | 0.030726293210986 |
| N0.HOG0001169 | OG0000985 | YcgN family cysteine cluster protein | 147 | 163 | 0.030451241122303 |
| N0.HOG0000814 | OG0000630 | rRNA maturation RNase YbeY | 149 | 172 | 0.026996962251038 |
| N0.HOG0000937 | OG0000753 | ribosome biogenesis factor YjgA | 148 | 208 | 0.02652326611546 |
| N0.HOG0001303 | OG0001119 | YigZ family protein | 140 | 250 | 0.025973897454513 |
| N0.HOG0001151 | OG0000967 | peroxide stress protein YaaA | 147 | 287 | 0.02181446243456 |
| N0.HOG0000763 | OG0000579 | DUF441 domain-containing protein | 148 | 153 | 0.021339223971957 |
| N0.HOG0000407 | OG0000226 | YicC/YloC family endoribonuclease | 150 | 294 | 0.020847593558978 |
| N0.HOG0001267 | OG0001083 | 30S ribosomal protein S12 methylthiotransferase accessory factor YcaO | 145 | 588 | 0.01880517000213 |
| N0.HOG0001290 | OG0001106 | YoaH family protein | 144 | 115 | 0.017301137594613 |
| N0.HOG0000761 | OG0000577 | YajQ family cyclic di-GMP-binding protein | 149 | 164 | 0.015026597633005 |
| N0.HOG0000795 | OG0000611 | HI1450 family dsDNA-mimic protein | 150 | 129 | 0.014993452268172 |
| N0.HOG0000991 | OG0000807 | tRNA (N6-isopentenyl adenosine(37)-C2)-methylthiotransferase MiaB | 148 | 478 | 0.0098674081507985 |
