## Supplementary material for "Type IV minor pilin ComN predicted the USS-receptor in Pasteurellaceae": Table S8.docx

**Table S8	Mean predicted probabilities for PpdA_Hin-USS_/PpdA_Apl-USS_ across all modeled species.** Predicted probabilities showing significant overrepresentation above random (0.25) are shown in bold and of those matching their respective USS in colour code (green A, blue C, orange G and red T.

| **Species and USS-clade** | **Mean probabilities** | | | | | **# input models** |
| --- | --- | --- | --- | --- | --- | --- |
|  | **A** | **C** | **G** | **T** | **USS** |  |
| ***Haemophilus influenzae* Rd**  **Hin-USS** | **0.4016***** | 0.1831*** | 0.2607 | 0.1546*** | **A** | **98** |
|  | **0.4727***** | 0.2536 | 0.1350*** | 0.1387*** | **A** |  |
|  | **0.3107***** | 0.1931*** | **0.2785*** | 0.2178*** | **G** |  |
|  | 0.0262*** | 0.2016*** | 0.0354*** | **0.7368***** | **T** |  |
|  | 0.1931*** | 0.1181*** | **0.3349***** | **0.3539***** | **G** |  |
|  | 0.0889*** | **0.5343***** | 0.0643*** | **0.3125***** | **C** |  |
|  | **0.4280***** | 0.0874*** | **0.4327***** | 0.0518*** | **G** |  |
|  | **0.3017***** | 0.1723*** | **0.4792***** | 0.0467*** | **G** |  |
|  | 0.1403*** | **0.4017***** | 0.1995*** | 0.2586 | **T** |  |
| ***Mannheimia succiniciproducens* strain MBEL55E**  **Hin-USS** | **0.6852***** | 0.0727*** | 0.1273*** | 0.1148*** | **A** | **90** |
|  | **0.4654***** | 0.1250*** | 0.1765*** | 0.2331 | **A** |  |
|  | 0.2169* | 0.1966*** | **0.3214**** | 0.2652 | **G** |  |
|  | 0.0380*** | 0.2749 | 0.1077*** | **0.5794***** | **T** |  |
|  | 0.2034** | 0.2423 | **0.3236*** | 0.2307 | **G** |  |
|  | 0.0771*** | **0.5687***** | 0.2354 | 0.1187*** | **C** |  |
|  | 0.2242 | 0.2564 | **0.4527***** | 0.0667*** | **G** |  |
|  | **0.3129***** | 0.2100* | **0.3820***** | 0.0951*** | **G** |  |
|  | 0.1439*** | 0.2478 | 0.1198*** | **0.4885***** | **T** |  |
| ***Aggregatibacter actinomycetemcomitans* strain 31S**  **Hin-USS** | **0.4307***** | 0.1583*** | 0.2674 | 0.1437*** | **A** | **47** |
|  | **0.3398***** | 0.2748 | 0.2315 | 0.1538*** | **A** | **4** |
|  | 0.1298*** | **0.4610***** | 0.2019** | 0.2073** | **G** |  |
|  | 0.0673*** | 0.2417 | 0.1298*** | **0.5612***** | **T** |  |
|  | 0.2121* | 0.1903** | 0.2861 | **0.3116*** | **G** |  |
|  | 0.2781 | 0.267 | 0.1089*** | **0.3460***** | **C** |  |
|  | **0.3216*** | 0.1342*** | 0.2676 | 0.2766 | **G** |  |
|  | **0.3063*** | 0.1471*** | **0.4678***** | 0.0787*** | **G** |  |
|  | 0.1745*** | **0.3540***** | 0.2501 | 0.2214* | **T** |  |
| ***Aggregatibacter sp* oral taxon 513**  **Hin-USS** | **0.3930***** | 0.2246* | 0.2476 | 0.1348*** | **A** | **85** |
|  | **0.2851*** | **0.3353***** | 0.2048*** | 0.1749*** | **A** |  |
|  | 0.1422*** | **0.3623***** | **0.2911*** | 0.2044*** | **G** |  |
|  | 0.0360*** | 0.1830*** | 0.0854*** | **0.6956***** | **T** |  |
|  | **0.3006*** | 0.1935*** | 0.27 | 0.2359 | **G** |  |
|  | 0.2467 | 0.2749 | 0.0652*** | **0.4132***** | **C** |  |
|  | **0.3680***** | 0.0753*** | **0.4494***** | 0.1073*** | **G** |  |
|  | **0.3662***** | 0.1526*** | **0.4034***** | 0.0778*** | **G** |  |
|  | 0.1689*** | **0.3716***** | 0.1839*** | **0.2756**** | **T** |  |
| ***Pasteurella multocida* strain NCTC8282**  **Hin-USS** | **0.3348***** | 0.2401 | **0.3112**** | 0.1139*** | **A** | **30** |
|  | 0.2777 | **0.3244**** | 0.2755 | 0.1223*** | **A** |  |
|  | 0.1997* | 0.2575 | **0.3879***** | 0.1549*** | **G** |  |
|  | 0.1069*** | 0.1984* | 0.1674*** | **0.5273***** | **T** |  |
|  | 0.2347 | 0.2407 | 0.2734 | 0.2511 | **G** |  |
|  | 0.1688* | 0.3106 | 0.0841*** | **0.4365***** | **C** |  |
|  | **0.3476**** | 0.234 | 0.2193 | 0.1991 | **G** |  |
|  | **0.5445***** | 0.0918*** | 0.2256 | 0.1381*** | **G** |  |
|  | 0.1109*** | 0.2575 | **0.2924*** | **0.3392***** | **T** |  |
| **All Hin-USS species** | **0.4706***** | 0.1663*** | 0.2285*** | 0.1346*** | **A** | **350** |
|  | **0.3907***** | 0.2493 | 0.1876*** | 0.1724*** | **A** |  |
|  | 0.2118*** | **0.2766**** | **0.2917***** | 0.2199*** | **G** |  |
|  | 0.0441*** | 0.2211** | 0.0901*** | **0.6448***** | **T** |  |
|  | 0.2279** | 0.1886*** | **0.3044***** | **0.2791**** | **G** |  |
|  | 0.1565*** | **0.4251***** | 0.1162*** | **0.3022***** | **C** |  |
|  | **0.3398***** | 0.1468*** | **0.4014***** | 0.1119*** | **G** |  |
|  | **0.3417***** | 0.1669*** | **0.4125***** | 0.0789*** | **G** |  |
|  | 0.1502*** | **0.3361***** | 0.1900*** | **0.3238***** | **T** |  |
| ***Actinobacillus equuli* subsp. haemolyticus strain 3524**  **Apl-USS** | **0.6227***** | 0.0722*** | 0.2415 | 0.0636*** | **A** | **96** |
|  | 0.0643*** | **0.4113***** | 0.0850*** | **0.4394***** | **C** |  |
|  | **0.5982***** | 0.0837*** | 0.1548*** | 0.1633*** | **A** |  |
|  | **0.4109***** | 0.1394*** | 0.1235*** | **0.3261***** | **A** |  |
|  | 0.1477*** | 0.2595 | 0.2109* | **0.3819***** | **G** |  |
|  | 0.0407*** | **0.3467***** | 0.1664*** | **0.4461***** | **C** |  |
|  | 0.1047*** | **0.3689***** | **0.4339***** | 0.0925*** | **G** |  |
|  | 0.223 | 0.2932 | **0.4285***** | 0.0553*** | **G** |  |
|  | 0.2097* | 0.2203 | **0.3830***** | 0.1869*** | **T** |  |
| ***Frederiksenia canicola* strain HPA 21**  **Apl-USS** | **0.5434***** | 0.1563*** | 0.1404*** | 0.1600*** | **A** | **76†** |
|  | 0.0360*** | **0.5299***** | 0.0331*** | **0.4010***** | **C** |  |
|  | **0.3649***** | 0.2252** | 0.0986*** | **0.3114***** | **A** |  |
|  | **0.5011***** | 0.1133*** | 0.0657*** | **0.3199***** | **A** |  |
|  | **0.3090***** | **0.3609***** | 0.1146*** | 0.2155*** | **G** |  |
|  | 0.1004*** | **0.3322***** | 0.0590*** | **0.5084***** | **C** |  |
|  | 0.1703*** | 0.1612*** | **0.5862***** | 0.0824*** | **G** |  |
|  | 0.1052*** | 0.1322*** | **0.7321***** | 0.0305*** | **G** |  |
|  | 0.1714*** | 0.0769*** | **0.5784***** | 0.1733*** | **T** |  |
| **Pasteurellaceae bacterium Orientalotternb1**  **Apl-USS** | 0.2108* | 0.2841* | **0.4037***** | 0.1014*** | **A** | **92** |
|  | 0.0311*** | **0.6032***** | 0.0867*** | 0.2789 | **C** |  |
|  | **0.4875***** | 0.1260*** | **0.2984*** | 0.0880*** | **A** |  |
|  | **0.4643***** | 0.1169*** | 0.1553*** | 0.2635 | **A** |  |
|  | **0.3275***** | 0.1578*** | 0.2176 | **0.2971**** | **G** |  |
|  | 0.1034*** | 0.2383 | 0.0751*** | **0.5832***** | **C** |  |
|  | 0.2616 | 0.1181*** | **0.4165***** | 0.2038* | **G** |  |
|  | **0.3063*** | 0.1712*** | **0.4337***** | 0.0887*** | **G** |  |
|  | 0.1358*** | **0.4056***** | 0.2247 | 0.234 | **T** |  |
| ***Mannheimia bovis* strain 39324S-11**  **Apl-USS** | 0.1581*** | 0.2459 | **0.5539***** | 0.0421*** | **A** | **98** |
|  | 0.0273*** | **0.6116***** | 0.0863*** | 0.2749 | **C** |  |
|  | **0.4388***** | 0.1564*** | 0.2721 | 0.1328*** | **A** |  |
|  | **0.4207***** | 0.0902*** | 0.1380*** | **0.3512***** | **A** |  |
|  | **0.2948**** | 0.1978*** | 0.2058*** | **0.3016**** | **G** |  |
|  | 0.1154*** | 0.2723 | 0.0657*** | **0.5467***** | **C** |  |
|  | **0.3070**** | 0.2055** | **0.4030***** | 0.0844*** | **G** |  |
|  | **0.4754***** | 0.1737*** | 0.2456 | 0.1053*** | **G** |  |
|  | 0.1555*** | 0.2355 | 0.2032*** | **0.4058***** | **T** |  |
| ***Actinobacillus lignieresii* strain NCTC4189**  **Apl-USS** | **0.3314***** | 0.1765*** | **0.4058***** | 0.0864*** | **A** | **96** |
|  | 0.0507*** | **0.4601***** | 0.0733*** | **0.4160***** | **C** |  |
|  | **0.5558***** | 0.1008*** | 0.2261 | 0.1173*** | **A** |  |
|  | **0.4042***** | 0.1678*** | 0.1509*** | 0.277 | **A** |  |
|  | 0.2281 | 0.2503 | 0.301 | 0.2206* | **G** |  |
|  | 0.0644*** | **0.4127***** | 0.0761*** | **0.4467***** | **C** |  |
|  | 0.2826 | 0.2868 | **0.3278**** | 0.1029*** | **G** |  |
|  | **0.4646***** | 0.0955*** | **0.2989*** | 0.1409*** | **G** |  |
|  | 0.0822*** | 0.2354 | **0.3581***** | **0.3242***** | **T** |  |
| **All Apl-USS species** | **0.3663***** | 0.1877*** | **0.3586***** | 0.0874*** | **A** | **458** |
|  | 0.0421*** | **0.5226***** | 0.0746*** | **0.3607***** | **C** |  |
|  | **0.4943***** | 0.1348*** | 0.2143*** | 0.1566*** | **A** |  |
|  | **0.4373***** | 0.1260*** | 0.1291*** | **0.3076***** | **A** |  |
|  | 0.2589 | 0.2408 | 0.2140*** | **0.2863***** | **G** |  |
|  | 0.0841*** | **0.3204***** | 0.0898*** | **0.5056***** | **C** |  |
|  | 0.2277* | 0.2319 | **0.4268***** | 0.1136*** | **G** |  |
|  | **0.3248***** | 0.1750*** | **0.4136***** | 0.0866*** | **G** |  |
|  | 0.1502*** | 0.2401 | 0.3399*** | **0.2697*** | **T** |  |

† Although 100/100 AF3 models had ipTM ≥ 0.6 and were used in DeepPBS, 24/100 of these were not used in the final DeepPBS output due to an unknown mismatch error in their DeepPBS predicted NumPy matrices.
