## Supplementary material for "Type IV minor pilin ComN predicted the USS-receptor in Pasteurellaceae": Table S9.docx

**Table S9	Mean predicted probabilities for PpdA_scr_ across all modeled species.** Predicted probabilities showing significant overrepresentation above random (0.25) are shown in bold and of those matching their respective USS in colour code (green A, blue C, orange G and red T.

| **Species and USS-clade** | **Mean probabilities** | | | | | **# input models** |
| --- | --- | --- | --- | --- | --- | --- |
|  | **A** | **C** | **G** | **T** | **USS** |  |
| ***Haemophilus influenzae* Rd**  **Hin-USS** | 0.2175* | **0.3437***** | 0.2044** | 0.2345 | **A** | **94** |
|  | 0.2778 | **0.3570**** | 0.1843*** | 0.1809*** | **A** |  |
|  | 0.2489 | 0.2497 | 0.2296 | 0.2718 | **G** |  |
|  | **0.2915*** | 0.1775*** | 0.2625 | 0.2685 | **T** |  |
|  | **0.3080**** | 0.2050** | 0.2218 | 0.2653 | **G** |  |
|  | **0.3070**** | 0.2030** | 0.2012** | 0.2887 | **C** |  |
|  | 0.2305 | 0.2327 | 0.2285 | **0.3082*** | **G** |  |
|  | 0.1799*** | 0.2291 | **0.3747***** | 0.2163 | **G** |  |
|  | 0.2048** | 0.2197 | **0.3482***** | 0.2273 | **T** |  |
| ***Mannheimia succiniciproducens* strain MBEL55E**  **Hin-USS** | **0.3310**** | 0.2054* | 0.2708 | 0.1927*** | **A** | **77** |
|  | 0.2285 | 0.1767*** | **0.3083*** | 0.2865 | **A** |  |
|  | 0.1806** | 0.1993* | **0.3409*** | 0.2793 | **G** |  |
|  | 0.2206 | 0.2950 | 0.2917 | 0.1928* | **T** |  |
|  | **0.3767***** | 0.2280 | 0.2207 | 0.1746*** | **G** |  |
|  | 0.2682 | 0.2519 | 0.2148 | 0.2652 | **C** |  |
|  | 0.2646 | 0.2596 | 0.2524 | 0.2234 | **G** |  |
|  | 0.2276 | 0.2934 | 0.1997* | 0.2793 | **G** |  |
|  | 0.1678*** | 0.2809 | 0.1774*** | **0.3739***** | **T** |  |
| ***Aggregatibacter actinomycetemcomitans* strain 31S**  **Hin-USS** | 0.2957 | 0.2899 | 0.2639 | 0.1504*** | **A** | **47** |
|  | 0.2772 | 0.2582 | 0.24 | 0.2245 | **A** |  |
|  | 0.2389 | 0.2464 | 0.2735 | 0.2412 | **G** |  |
|  | **0.3119*** | 0.1920** | 0.1896** | 0.3066 | **T** |  |
|  | 0.2056* | 0.2513 | 0.2009* | **0.3422**** | **G** |  |
|  | 0.2476 | 0.2167 | 0.2114 | **0.3243**** | **C** |  |
|  | 0.2212 | 0.2497 | 0.3128 | 0.2163 | **G** |  |
|  | 0.2346 | 0.2747 | 0.2801 | 0.2107* | **G** |  |
|  | 0.1883** | 0.2741 | **0.3060**** | 0.2315 | **T** |  |
| ***Aggregatibacter sp* oral taxon 513**  **Hin-USS** | 0.2533 | **0.3298***** | 0.2067** | 0.2101* | **A** | **82** |
|  | 0.2485 | 0.2581 | 0.2487 | 0.2447 | **A** |  |
|  | 0.2678 | **0.3279**** | 0.1743*** | 0.23 | **G** |  |
|  | **0.3102*** | 0.2311 | 0.1943** | 0.2644 | **T** |  |
|  | 0.2561 | 0.2231 | 0.2194 | **0.3014*** | **G** |  |
|  | 0.2376 | 0.2194 | 0.2117 | **0.3313***** | **C** |  |
|  | 0.2611 | 0.2145 | 0.2885 | 0.2359 | **G** |  |
|  | 0.2629 | 0.2077 | **0.3180*** | 0.2114 | **G** |  |
|  | 0.1766*** | 0.2299 | **0.3577***** | 0.2358 | **T** |  |
| ***Pasteurella multocida* strain NCTC8282**  **Hin-USS** | 0.2268 | **0.2886*** | 0.2601 | 0.2245 | **A** | **66** |
|  | 0.258 | 0.2516 | **0.3302**** | 0.1601*** | **A** |  |
|  | 0.2637 | **0.3028*** | 0.2123* | 0.2212 | **G** |  |
|  | 0.2742 | 0.2055 | 0.2219 | **0.2985*** | **T** |  |
|  | **0.3234**** | 0.1614*** | 0.2416 | 0.2736 | **G** |  |
|  | 0.2783 | 0.2036** | 0.2224 | 0.2957 | **C** |  |
|  | 0.2284 | 0.2291 | 0.2658 | 0.2768 | **G** |  |
|  | 0.2205 | 0.263 | **0.2994*** | 0.2171 | **G** |  |
|  | 0.1709*** | **0.3371***** | 0.245 | 0.247 | **T** |  |
| **All Hin-USS species** | 0.2611 | 0.2946*** | 0.2366 | 0.2076*** | **A** | **366** |
|  | 0.2572 | 0.2652 | 0.2583 | 0.2192** | **A** |  |
|  | 0.2402 | 0.2658 | 0.2431 | 0.2509 | **G** |  |
|  | 0.2803** | 0.2211* | 0.2367 | 0.2619 | **T** |  |
|  | 0.3005*** | 0.2120*** | 0.2219** | 0.2657 | **G** |  |
|  | 0.2705* | 0.2188*** | 0.2115*** | 0.2991*** | **C** |  |
|  | 0.243 | 0.2358 | 0.2645 | 0.2567 | **G** |  |
|  | 0.2229** | 0.2498 | **0.2994***** | 0.2279* | **G** |  |
|  | 0.1825*** | 0.263 | 0.2904*** | 0.2641 | **T** |  |
| ***Actinobacillus equuli* subsp. haemolyticus strain 3524**  **Apl-USS** | 0.2497 | 0.2165 | **0.3791***** | 0.1547*** | **A** | **80** |
|  | 0.1616*** | 0.2411 | **0.3825***** | 0.2148 | **C** |  |
|  | 0.2681 | 0.1946** | 0.2547 | 0.2826 | **A** |  |
|  | **0.3137**** | 0.2025* | 0.1983* | 0.2855 | **A** |  |
|  | **0.3588***** | 0.1776*** | 0.1390*** | **0.3245**** | **G** |  |
|  | **0.3106*** | 0.2079* | 0.2000** | 0.2815 | **C** |  |
|  | 0.2785 | 0.2764 | 0.2167 | 0.2284 | **G** |  |
|  | 0.2363 | **0.3489**** | 0.2335 | 0.1813*** | **G** |  |
|  | 0.1207*** | **0.3479***** | 0.2548 | 0.2767 | **T** |  |
| ***Frederiksenia canicola* strain HPA 21**  **Apl-USS** | 0.2048* | **0.3183**** | 0.2941 | 0.1829*** | **A** | **91** |
|  | 0.2326 | 0.2916 | 0.2853 | 0.1905*** | **C** |  |
|  | 0.2585 | 0.2498 | 0.2254 | 0.2663 | **A** |  |
|  | 0.2893 | 0.2137 | 0.1374*** | **0.3596***** | **A** |  |
|  | **0.3617***** | 0.1804*** | 0.1496*** | **0.3083*** | **G** |  |
|  | **0.3644***** | 0.1784*** | 0.1752*** | 0.2821 | **C** |  |
|  | 0.2464 | 0.2482 | 0.2769 | 0.2285 | **G** |  |
|  | 0.2054* | 0.2825 | **0.3236**** | 0.1885*** | **G** |  |
|  | 0.1711*** | 0.2513 | **0.3720***** | 0.2056* | **T** |  |
| **Pasteurellaceae bacterium Orientalotternb1**  **Apl-USS** | 0.1797*** | **0.3225***** | **0.2982*** | 0.1995*** | **A** | **88** |
|  | 0.2238 | 0.2969 | 0.2862 | 0.1932** | **C** |  |
|  | 0.2469 | 0.2885 | 0.2175 | 0.2472 | **A** |  |
|  | **0.3407***** | 0.2063* | 0.1968** | 0.2561 | **A** |  |
|  | **0.4048***** | 0.1610*** | 0.1648*** | 0.2694 | **G** |  |
|  | 0.3001 | 0.1793** | 0.1962* | **0.3245**** | **C** |  |
|  | 0.2851 | 0.1729*** | **0.3076*** | 0.2344 | **G** |  |
|  | 0.2271 | 0.282 | 0.2855 | 0.2054* | **G** |  |
|  | 0.1561*** | **0.3133***** | 0.2753 | 0.2553 | **T** |  |
| ***Mannheimia bovis* strain 39324S-11**  **Apl-USS** | 0.2483 | 0.2335 | **0.3797***** | 0.1386*** | **A** | **76** |
|  | 0.2396 | 0.26 | 0.2103* | 0.2901 | **C** |  |
|  | 0.2239 | 0.2965 | 0.242 | 0.2376 | **A** |  |
|  | 0.22 | 0.1689*** | 0.2063 | **0.4048***** | **A** |  |
|  | 0.2864 | 0.2077* | 0.1789*** | **0.3269***** | **G** |  |
|  | **0.3568***** | 0.2019 | 0.2393 | 0.2020* | **C** |  |
|  | **0.3438**** | 0.2645 | 0.2541 | 0.1376*** | **G** |  |
|  | 0.241 | 0.248 | 0.2662 | 0.2448 | **G** |  |
|  | 0.1288*** | **0.3361***** | 0.2469 | 0.2882 | **T** |  |
| ***Actinobacillus lignieresii* strain NCTC4189**  **Apl-USS** | 0.2618 | 0.2268 | **0.2988*** | 0.2127* | **A** | **96** |
|  | 0.2823 | 0.2065* | 0.2841 | 0.227 | **C** |  |
|  | 0.1671*** | **0.3443***** | 0.2111* | 0.2774 | **A** |  |
|  | 0.2755 | 0.2528 | 0.2495 | 0.2223 | **A** |  |
|  | 0.1974* | 0.2734 | 0.1921* | **0.3371**** | **G** |  |
|  | **0.4152***** | 0.1492*** | **0.3161*** | 0.1196*** | **C** |  |
|  | **0.3122**** | 0.2375 | 0.2242 | 0.226 | **G** |  |
|  | 0.2644 | 0.2665 | 0.1862*** | 0.2828 | **G** |  |
|  | 0.1767*** | 0.2592 | 0.2832 | 0.2809 | **T** |  |
| **All Apl-USS species** | 0.2284* | 0.2649 | **0.3268***** | 0.1799*** | **A** | **431** |
|  | 0.2299* | 0.2588 | **0.2900***** | 0.2213** | **C** |  |
|  | 0.2315* | **0.2767*** | 0.2290* | 0.2628 | **A** |  |
|  | **0.2890***** | 0.2109*** | 0.1980*** | **0.3021***** | **A** |  |
|  | **0.3201***** | 0.2014*** | 0.1654*** | **0.3131***** | **G** |  |
|  | **0.3512***** | 0.1817*** | 0.2268* | **0.2403** | **C** |  |
|  | **0.2921***** | 0.2386 | 0.2562 | 0.2131*** | **G** |  |
|  | 0.235 | **0.2851**** | 0.2584 | 0.2216** | **G** |  |
|  | 0.1525*** | **0.2986***** | **0.2887***** | 0.2603 | **T** |  |
